# coMethDMR: Accurate identification of co-methylated and differentially methylated regions in epigenome-wide association studies

**DOI:** 10.1101/615427

**Authors:** Lissette Gomez, Gabriel J. Odom, Juan I. Young, Eden R. Martin, Lizhong Liu, Xi Chen, Anthony J. Griswold, Zhen Gao, Lanyu Zhang, Lily Wang

## Abstract

Recent technology has made it possible to measure DNA methylation profiles in a cost-effective and comprehensive genome-wide manner using array-based technology for epigenome-wide association studies. However, identifying differentially methylated regions (DMRs) remains a challenging task because of the complexities in DNA methylation data. Supervised methods typically focus on the regions that contain consecutive highly significantly differentially methylated CpGs in the genome, but may lack power for detecting small but consistent changes when few CpGs pass stringent significance threshold after multiple comparison. Unsupervised methods group CpGs based on genomic annotations first and then test them against phenotype, but may lack specificity because the regional boundaries of methylation are often not well defined. We present `coMethDMR`, a flexible, powerful, and accurate tool for identifying DMRs. Instead of testing all CpGs within a genomic region, `coMethDMR` carries out an additional step that selects co-methylated sub-regions first. Next, `coMethDMR` tests association between methylation levels within the sub-region and phenotype via a random coefficient mixed effects model that models both variations between CpG sites within the region and differential methylation simultaneously. `coMethDMR` offers well-controlled Type I error rate, improved specificity, focused testing of targeted genomic regions, and is available as an open-source R package.

## INTRODUCTION

Many diseases are caused by a complex interplay of genes and environment factors, such as smoking, poor diet, and lack of exercise. Epigenetic studies investigate the mechanisms that modify the expression levels of selected genes without changes to the underlying DNA sequence. The study of these epigenetic patterns hold excellent promise for detecting new regulatory mechanisms that may be susceptible to modification by environmental factors, which in turn increase the risk of disease. Among epigenetic modifications, DNA methylation is the most widely studied. The addition or removal of a methyl group at the 5th position of a cytosine is the key feature of DNA methylation. Alterations of DNA methylation levels have been shown to be involved in many diseases (1), such as cancers (2–4) and neurodegenerative diseases (5–7).

While whole-genome bisulfite sequencing is still too costly for large epidemiologic studies, recent technology has made it possible to measure DNA methylation profiles in a cost-effective and comprehensive genome-wide manner using array-based technology such as the Infinium MethylationEPIC BeadChip Kit (8), which allows researchers to interrogate more than 850,000 methylation sites per sample at single-nucleotide resolution. The first wave of epigenome analysis tools have focused on comprehensive DNA methylation analysis of single base sites, that is, identifying differentially methylated CpG sites, while more recent effort have shifted to analyzing differentially methylated regions (DMRs) (9–11).

The shift to DMR analyses are driven by both biological and statistical reasons. Biologically, it has been observed that methylation levels are strongly correlated across the genome and methylation often occurs as a regional phenomenon (12). In the study of complex diseases, various studies have reported functionally-relevant genomic regions, such as CpG islands (13) or CpG island shores (14), are associated with diseases. While changes at single sites should not be overlooked, DMRs have increasingly been deemed the hallmarks of differential methylation and replication of DMRs are often more successful compared with changes at single sites (15,16). Statistically, because of the large number of CpG sites interrogated by methylation arrays, testing regions rather than individual CpGs can help improve power by reducing the number of tests conducted. In addition, while effect size in a single CpG might be small, by borrowing information from all the CpGs within a region, statistical test for regions can more effectively leverage information within the region to increase sensitivity and specificity.

A number of statistical methods for identifying DMRs have been proposed (10,11,17–20), reviewed (21,22), and compared (23–25). Methods for DMR identification can be classified into supervised and unsupervised methods. Supervised methods (e.g. `bumphunter` (17), `DMRcate` (10), and `probeLasso` (11)) typically start with computing p-values for differential methylation at individual CpG sites, and then scan the genome to identify regions with adjacent low p-values. The statistical significance of these regions is then computed by combining individual CpG p-values in the region using methods such as Stouffer’s Z (10). However, a challenge with supervised methods is that they may lack power for detecting small but consistent changes when few CpGs pass stringent significance threshold after multiple comparison (25).

An alternative strategy is to use an unsupervised approach, which defines relevant regions across the genome first, independently of any phenotype information, and then tests methylation levels in these predefined regions against a phenotype (19,20,26). In this study, we propose a new unsupervised approach for testing differential methylation in regions against continuous phenotype such as age.

Table 1 lists several previously proposed unsupervised methods. A challenge with unsupervised approaches is their lack of specificity. Unlike gene expression data, the regional boundary of DNA methylation is often not well defined. Therefore, currently available approaches that summarize methylation levels in a region using mean or median methylation levels of the CpGs within the region may have results that vary depending on the boundaries of the region. In addition, when testing associations between phenotype and the summarized methylation levels in a genomic region, the spatial correlations between CpG sites within the region is ignored.

**Table 1.**
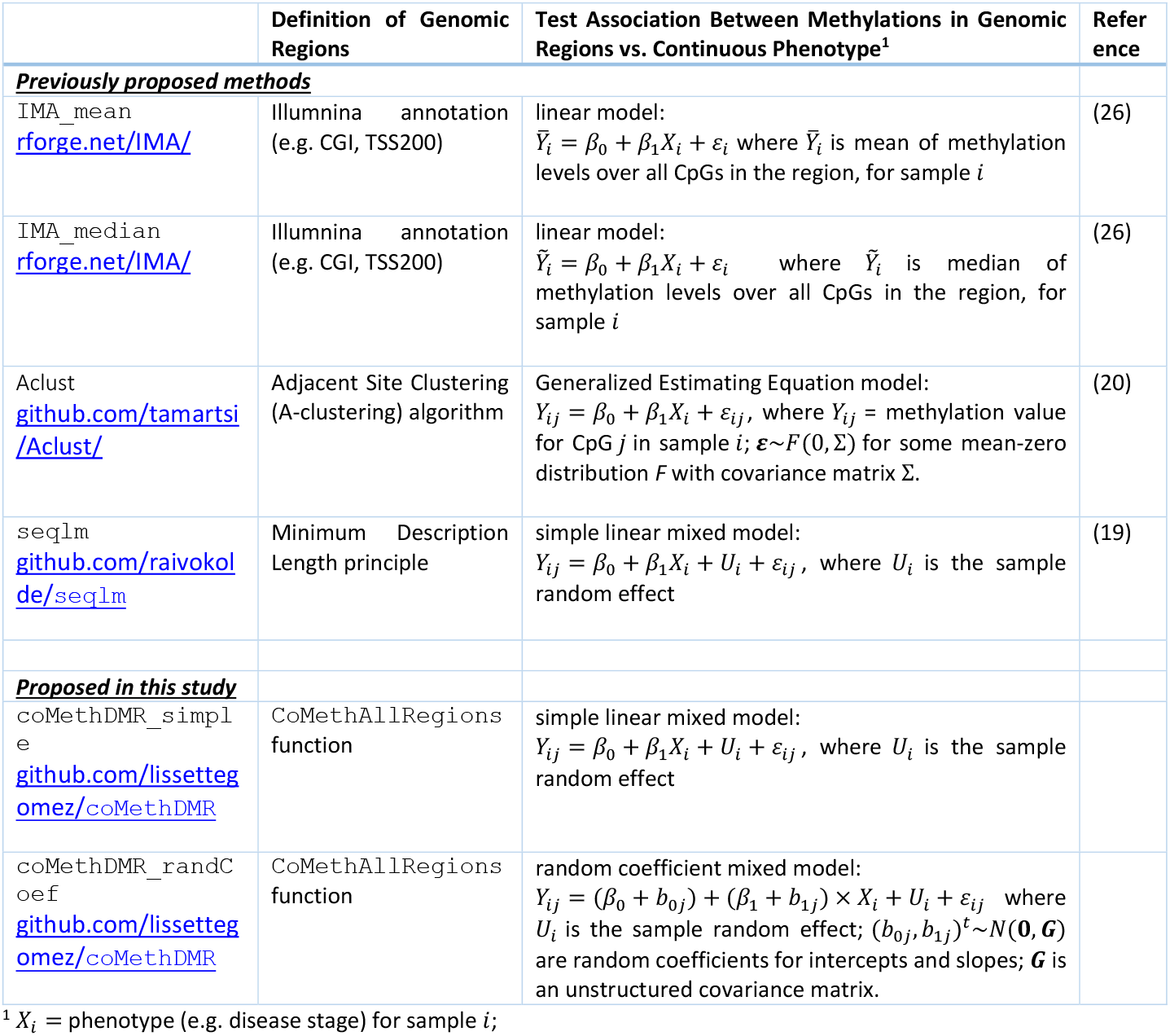
Summary of unsupervised methods

Here we present `coMethDMR`, a new unsupervised approach that optimally leverages covariations among CpGs within genomic regions to identify genomic regions associated with continuous phenotypes. Instead of testing all CpGs within a genomic region, `coMethDMR` carries out an additional step that clusters co-methylated sub-regions without using any outcome information. Next, `coMethDMR` tests association between methylation within the sub-region and continuous phenotype using a random coefficient mixed effects model (27), which models both variations between CpG sites within the region and differential methylation simultaneously.

In the following sections, we provide methodological details of the `coMethDMR` analysis pipeline and compare this new method with several existing tools. The advantages of `coMethDMR` is demonstrated using both simulated and real methylation datasets. We show that the additional CpG selection step (subregion identification) improves power substantially while preserving Type I error rate. In addition, the new random coefficient model improves specificity and is robust against association signals from outlier CpGs when detecting changes in differential methylation in the regions.

## MATERIAL AND METHODS

### The coMethDMR analysis pipeline

Figure 1 shows the workflow of the `coMethDMR` analysis pipeline. There are two major steps in the `coMethDMR` pipeline: (1) within a genomic region, identify the sub-region with contiguous and co-methylated CpGs, and (2) test association of CpG methylation in the sub-region with phenotype, while modelling for variabilities among the CpGs simultaneously.

**Figure 1.**
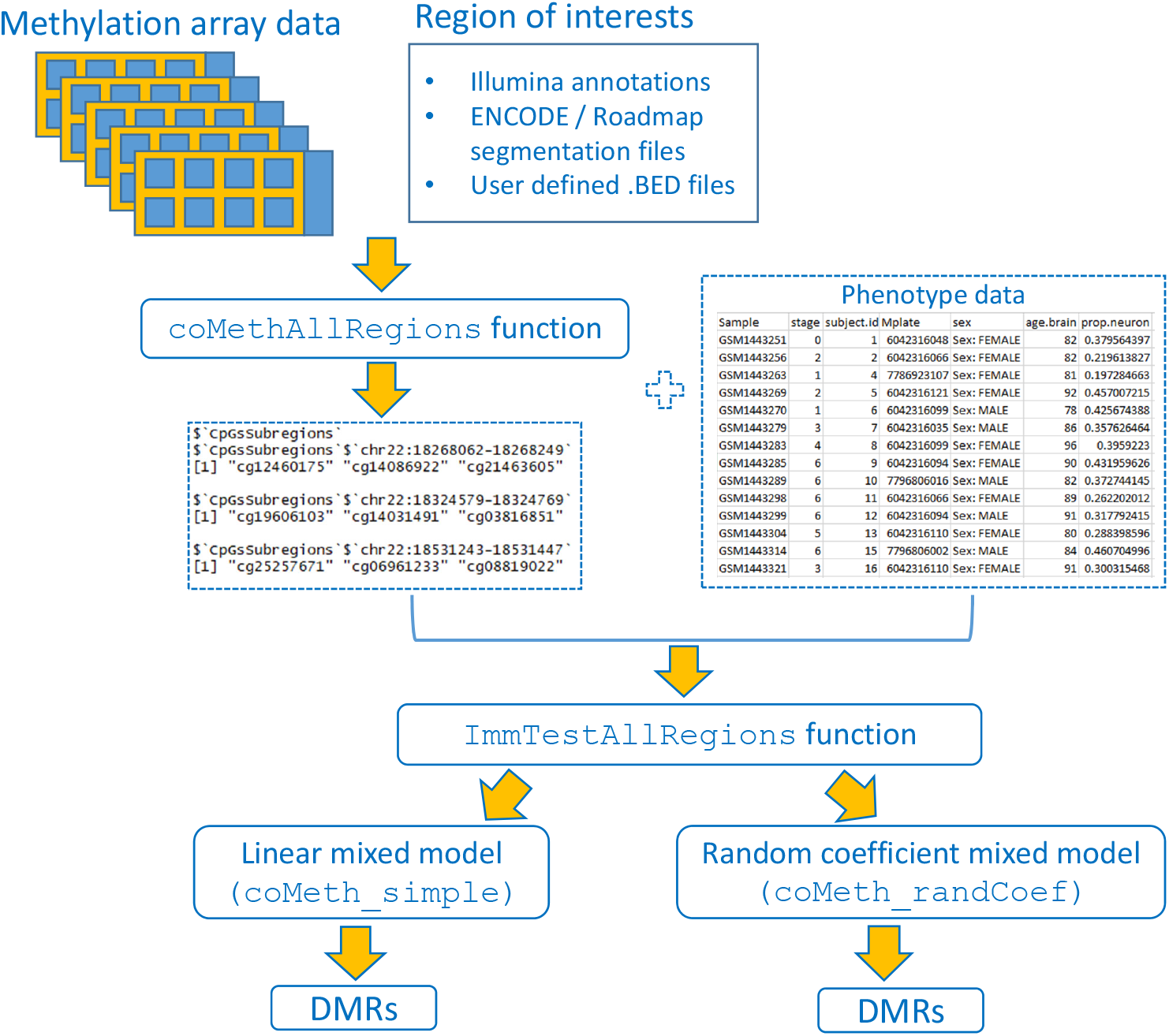
Workflow of the `coMethDMR` analysis pipeline.

In the first step, the genome will be divided into regions by taking advantage of methylation array annotations. Because the Illumina chips target methylation sites primarily at genic regions and CpG islands (CGIs, regions in the genome where there are more CG dinucleotides found than expected by chance), the regions can be defined based on their relations to genes or CGIs. Figure 2A shows correlation between methylation levels among CpGs in an example of a genomic region corresponding to the CGI located at `chr10:100028236-100028499`. This region includes 7 CpG probes ordered by their locations on the chromosome. Note that in spite of belonging to the same CGI, only the last 4 probes constitute the co-methylated region. To select the co-methylated region, we use the rdrop statistic, which is the correlation between each CpG with the sum of methylation levels in all other CpGs (Fig 2B). Note that in this example, the co-methylated region consists of all the contiguous CpGs with rdrop statistics greater than 0.5. We evaluated the sensitivity and specificity of the rdrop statistic at identifying co-methylated CpGs in the subsection “Optimal parameter for in `coMethDMR` pipeline” below.

**Figure 2.**
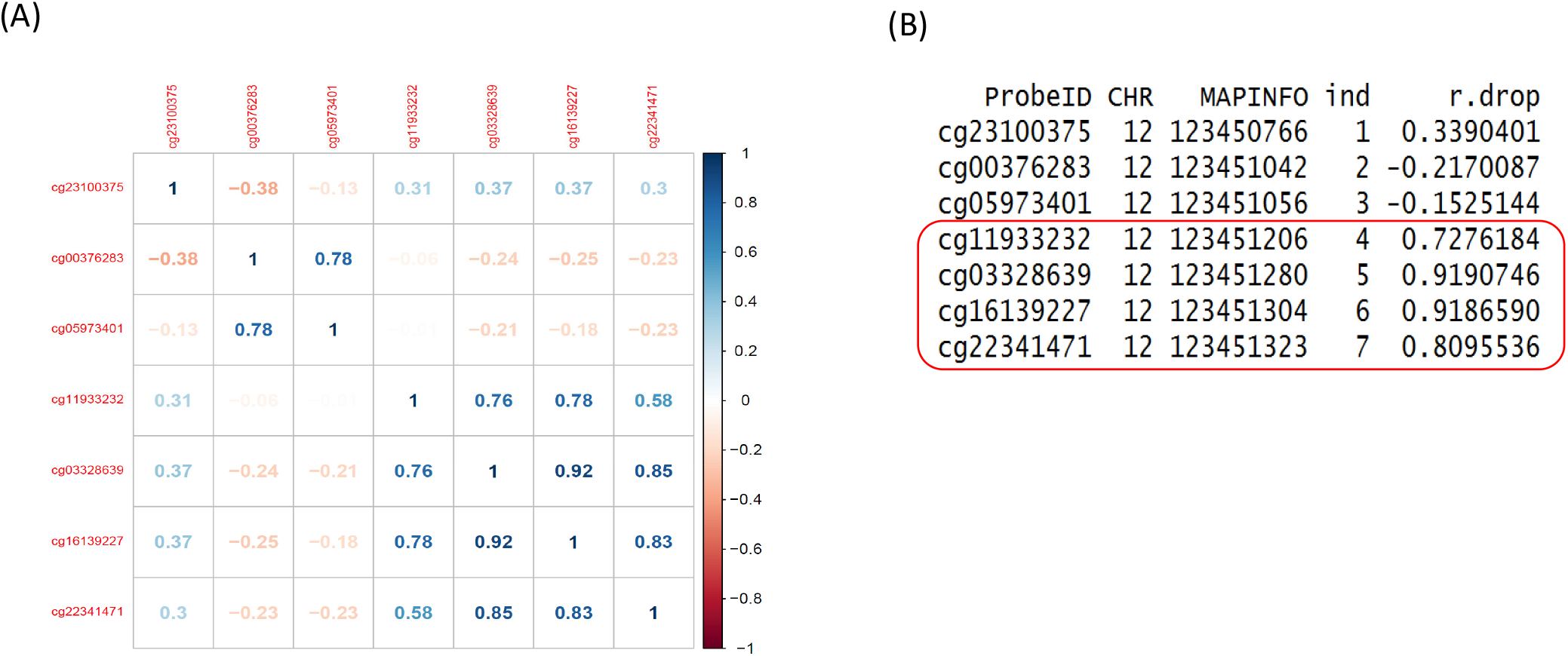
An example of a contiguous co-methylated sub-region. (A) This pre-defined region (a CGI) included 7 CpG probes ordered by their location. Shown are correlation between methylation levels in each pair of CpGs. Note that only the last 4 probes constitute the co-methylated region within this CGI. (B) The CpGs in the co-methylated sub-region can be identified using the rdrop statistic, which is the correlation between each CpG with the sum of methylation levels in all other CpGs. In this example, all the co-methylated CpGs had rdrop statistics greater than 0.5.

In the second step, to simultaneously model variations among the co-methylated CpGs as well as association with phenotype, we propose a random coefficient mixed model for testing groups of CpGs against phenotype. Figure 3 provides a hypothetical example fit of the mixed model for testing two CpGs against disease stage (treated as a linear variable). This model includes (1) normalized methylation values as the outcome variable, (2) a systematic component that models the mean for each group of CpGs (the fixed effects intercept *β*_0_ and slope *β*_1_ for variable stage), and (3) a random component (the random coefficients) that model how each CpG’s slope for stage varies about the group mean (the random effects *b*_0*j*_ and *b*_1*j*,_*j* = 1,2). Because both fixed and random effects are included in this model, this model is a mixed effects model. Additional details of the random coefficient model are described in Supplementary Text.

**Figure 3.**
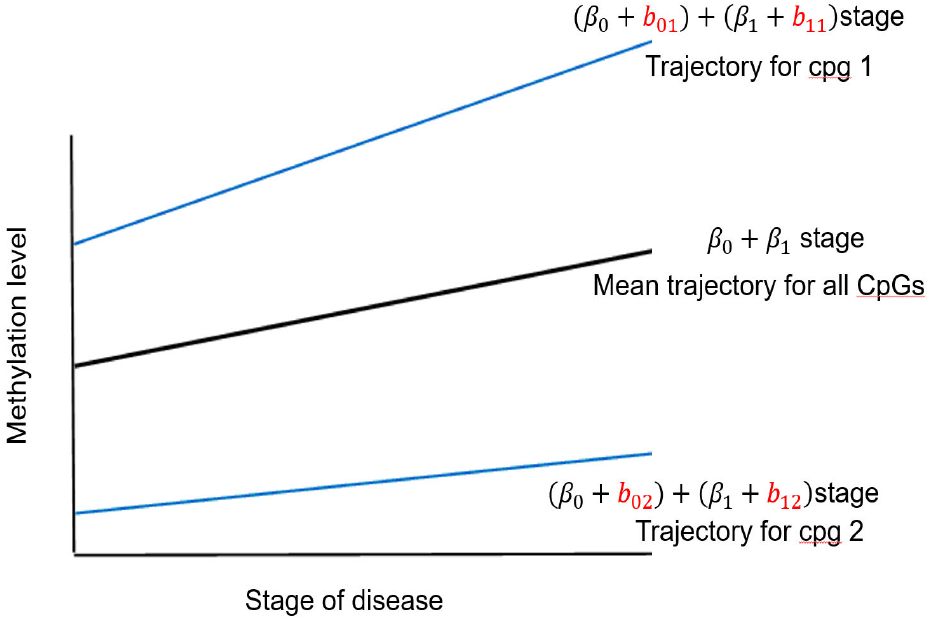
Illustration – proposed random coefficient mixed model for testing methylation levels in a hypothetical genomic region with 2 CpGs against disease stage treated as a linear variable.

We are interested in testing the null hypothesis that there is no association between phenotype (disease stage) and methylation values. This can be accomplished by testing the fixed effect for slope *H*_0_:*β*_1_ = 0. In the sections below, we compare the statistical properties (i.e. power and Type I error rate) of this new random coefficient model with several currently available statistical models shown in Table 1.

## RESULTS

### Optimal parameter for in coMethDMR pipeline

The only parameter in the entire `coMethDMR` pipeline is the `rdrop` threshold in the identification of co-methylated sub-regions within a genomic region (Figure 2B). These rdrop statistics are the leave-one-out correlations between each CpG with the sum of methylation levels in all other CpGs. The co-methylated regions can be identified by ordering the CpGs by location, and selecting contiguous CpGs with leave-one-out correlations greater than a pre-specified threshold (`rdrop`), such as 0.5.

#### Simulation study 1

We conducted an analysis to assess the sensitivity and specificity of different `rDrop` values at identifying co-methylated CpGs. For each of the 19977 CGI regions with at least 3 CpGs, we computed pairwise correlations of the CpGs within each CGI region. Next, we selected regions with 3, 5, or 8 CpGs (parameter `ncpgs`) that have average pairwise correlations between 0.5-0.8 or 0.8 to 1 (parameter `minCorr`) for this simulation study. For each genomic region, we added additional irrelevant CpGs by sampling CpGs randomly from the genome. The number of random CpGs added were either the same as ncpgs (parameter `fold` = 1), or two times of ncpgs (parameter `fold` = 2). Therefore, by design of the experiment, the status of each CpG, i.e. whether they belong to a co-methylated cluster or not, is known.

These parameter values yielded 12 simulation scenarios: `ncpgs` (3, 5, or 8) x `minCorr` (0.5-0.8 or 0.8-1) x `fold` (1 or 2). For each scenario, this process was repeated 10 times to generate a total of 120 simulation datasets. Step 1 in the `coMethDMR` pipeline was used to identify co-methylated regions for each simulated genomic region. Supplementary Table 1 and Figures 4-5 show average sensitivity, specificity, and area under the ROC curve (AUC) for each value of the rDrop parameter. Optimal AUC occurred at or near `rDrop` = 0.4 for most of the simulation scenarios (Supplementary Table 1). Figure 4 and 5 also showed that at `rDrop` = 0.4, both sensitivity and specificity were over 0.9 for all but one simulation scenario. Therefore, we set `rDrop` at 0.4 for subsequent analysis.

**Figure 4.**
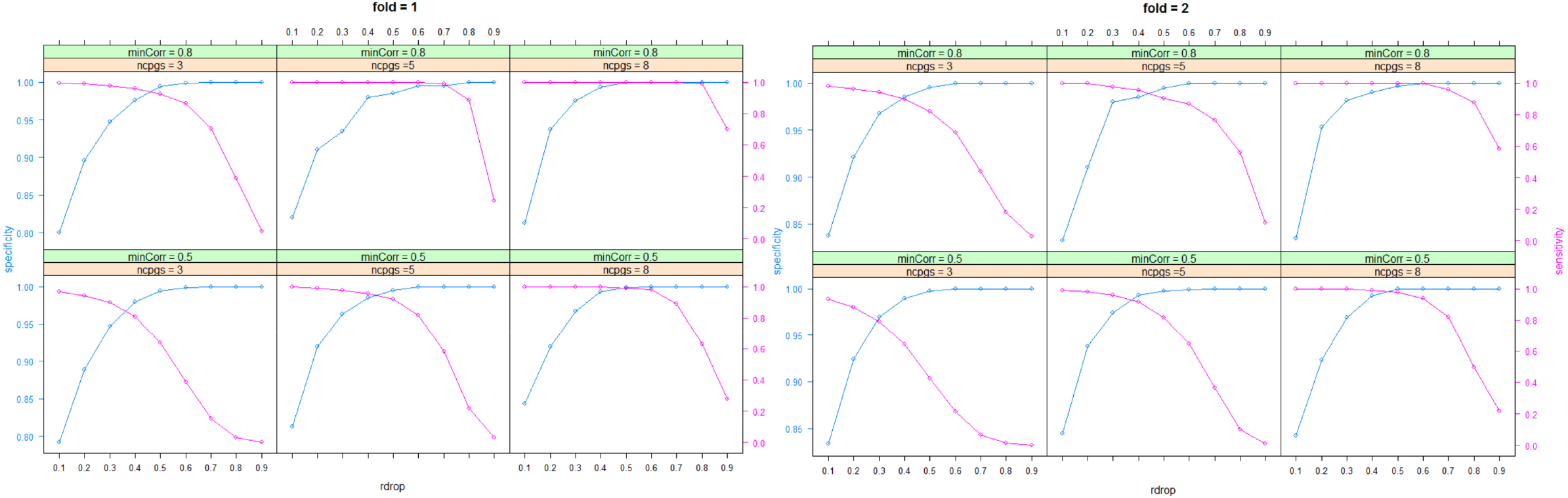
Optimal sensitivities and specificities for `coMethDMR` were achieved when the parameter `rDrop` is close to 0.4.

**Figure 5.**
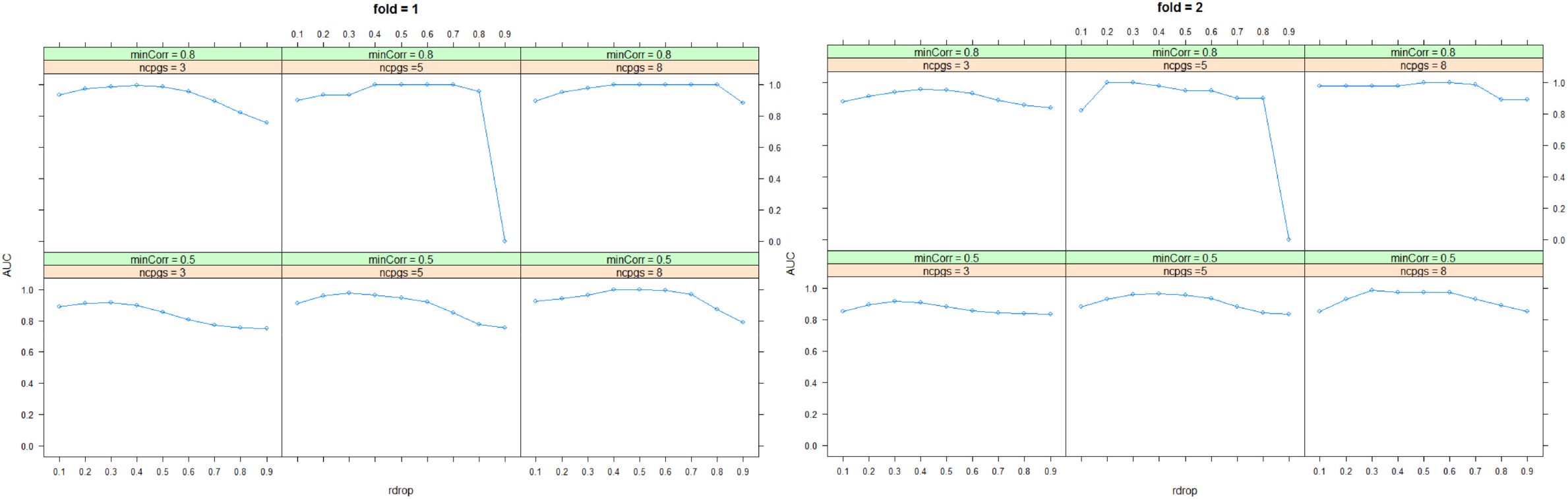
Optimal area under ROC curve (AUC) for `coMethDMR` was also achieved when the parameter `rDrop` is close to 0.4.

### coMethDMR controls Type I error when testing association between methylation levels in the genomic regions with phenotype

We conducted two simulation studies to assess the Type I error (i.e. false positive rate) of the proposed method. In <u>*Simulation study 2*</u>, we assume the genomic regions are pre-determined and we compared results using different statistical models for testing association of methylation levels with randomly generated phenotypes. In <u>*Simulation study 3*</u>, we assume the genomic regions are not pre-determined, and we determined Type I error for the entire `coMethDMR` pipeline. That is, we first identified co-methylated regions, then tested these selected regions for association with randomly generated phenotypes.

#### Simulation study 2

We evaluated the Type I error rate of several currently available statistical methods, along with the newly-proposed random coefficient mixed model, by generating random phenotype data and test their association with genomic regions in a real DNA methylation dataset. More specifically, we compared the five statistical models listed in Table 1: (1) a linear model with mean methylation *M* values as summary for a genomic region, (2) a linear model with median *M* values as summary for a genomic region, (3) a GEE model, (4) a simple linear mixed model, and (5) our proposed random coefficient linear mixed model.

To emulate correlation structure between different CpGs in real data, we generated simulation datasets using a real methylation dataset (GSE59685) as input. Lunnon et al. (2014) conducted an AD study that measured DNA methylation levels in four brain regions postmortem from 122 individuals using the Infinium HumanMethylation450K BeadChip platform (7). For this Type I error analysis, we used prefrontal cortex methylation data from 27 control subjects. For each simulation dataset, we randomly generated an age value for each sample. In the following sections, we use the term `pseudo age` to refer to the computer-simulated age variable.

First, we select 10 CpG island genomic regions randomly. For each region, we generated `pseudo age` randomly from a Poisson distribution with mean 65, independently of the methylation data. Therefore, by design of experiment, these `pseudo age` values are not associated with any methylation regions. This procedure was repeated 1000 times, to generate 10,000 simulation datasets (10 genomic regions x 1000 repetitions). Under the null hypothesis of no association between methylation and `pseudo age`, we expected the *p*-value distributions for a model to follow the uniform distribution, where 5% of the *p*-values would be less than 0.05, corresponding to a Type I error rate of 0.05.

In Figure 6, the estimated Type I error rates were 0.1011 (GEE model), 0.0545 (linear model with mean summary), 0.0532 (linear model with median summary), 0.0404 (random coefficient mixed model) and 0.0002 (simple linear mixed model). We can see that the GEE model showed inflated false positive rate, with highest Type I error at around 0.1. The inflated type I error rate by GEE model was also observed recently in simulation studies conducted by another group (19). On the other hand, the simple linear mixed model in (4) was overly conservative, with Type I error around 0.0002. Among the models, the proposed random coefficient mixed model in (5) and the linear models with mean or median summary had type I error closest to 0.05.

**Figure 6.**
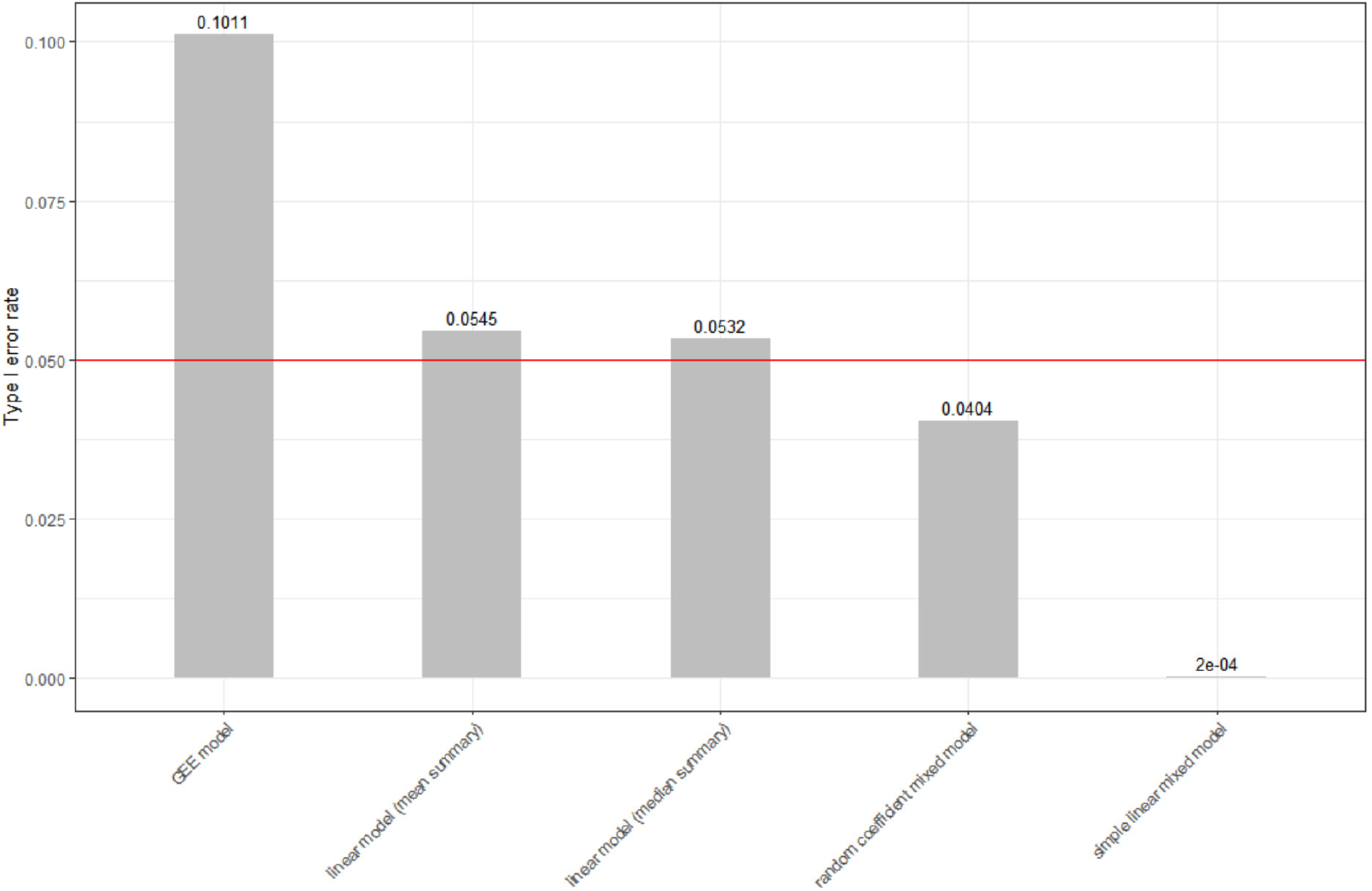
Type I error rates in the absence of differential methylation for different statistical models. Shown above the bars are proportions of CpG island genomic regions with p-values less than 0.05, for association with randomly generated “age” from Poisson distribution with mean 65, average over 10,000 simulation datasets. Under the null hypothesis of no association, we expected p-values to follow a uniform distribution, so methods that control type I error at nominal level would be close to the red line.

#### Simulation study 3

We next determined Type I error rate for `coMethDMR` when genomic regions are not pre-determined. That is, we determined if the `coMethDMR` pipeline, which includes both identifying co-methylated methylation clusters, and testing methylation regions against phenotype using linear mixed models, would still have controlled Type I error rates. To this end, we first identified 4444 co-methylated genomic regions mapped to CpG islands. Next we selected 10 co-methylated regions randomly, and then repeated <u>*Simulation study 2*</u> on these co-methylated regions. That is, for each co-methylated region, `pseudo age` values were generated randomly from Poisson distribution with mean of 65 for each of the samples. This procedure was repeated 1000 times to generate 10,000 simulation datasets (10 co-methylated regions x 1000 repetitions). Suppl. Figure 1 shows that after selecting co-methylated CpGs, Type I error remained well-controlled for both mixed models.

### coMethDMR improves power substantially compared to fitting mixed model directly to methylation data

#### Simulation study 4

Because genomic region are typically defined *a priori* based on annotations, without regard to the methylation data sets to be analyzed, we expect only a subset of CpGs in a pre-defined genomic region would be associated with the phenotype. We hypothesized that power can be improved by selecting consecutive CpGs in the co-methylated region first. To assess the power of the models that had controlled Type I error rate (i.e. simple linear mixed model and random coefficient mixed model), we performed a simulation study similar to <u>*Simulation study 2*</u> described above, except by testing methylation levels in the genomic regions against randomly generated `pseudo age` that are ranked in the same order as the mean of methylation values in the co-methylated sub-region. Therefore, by design of the experiment, the values of `pseudo ages` of the samples are associated with co-methylated CpGs in each genomic region.

The results in Figure 7 show that for both mixed models, power improved substantially after selecting the co-methylated regions. Without selecting co-methylated CpGs, the random coefficient mixed model has more power. After selecting co-methylated CpGs, both models performed similarly, especially when the number of co-methylated CpGs was at least moderate (i.e. nCpGs >= 5).

**Figure 7.**
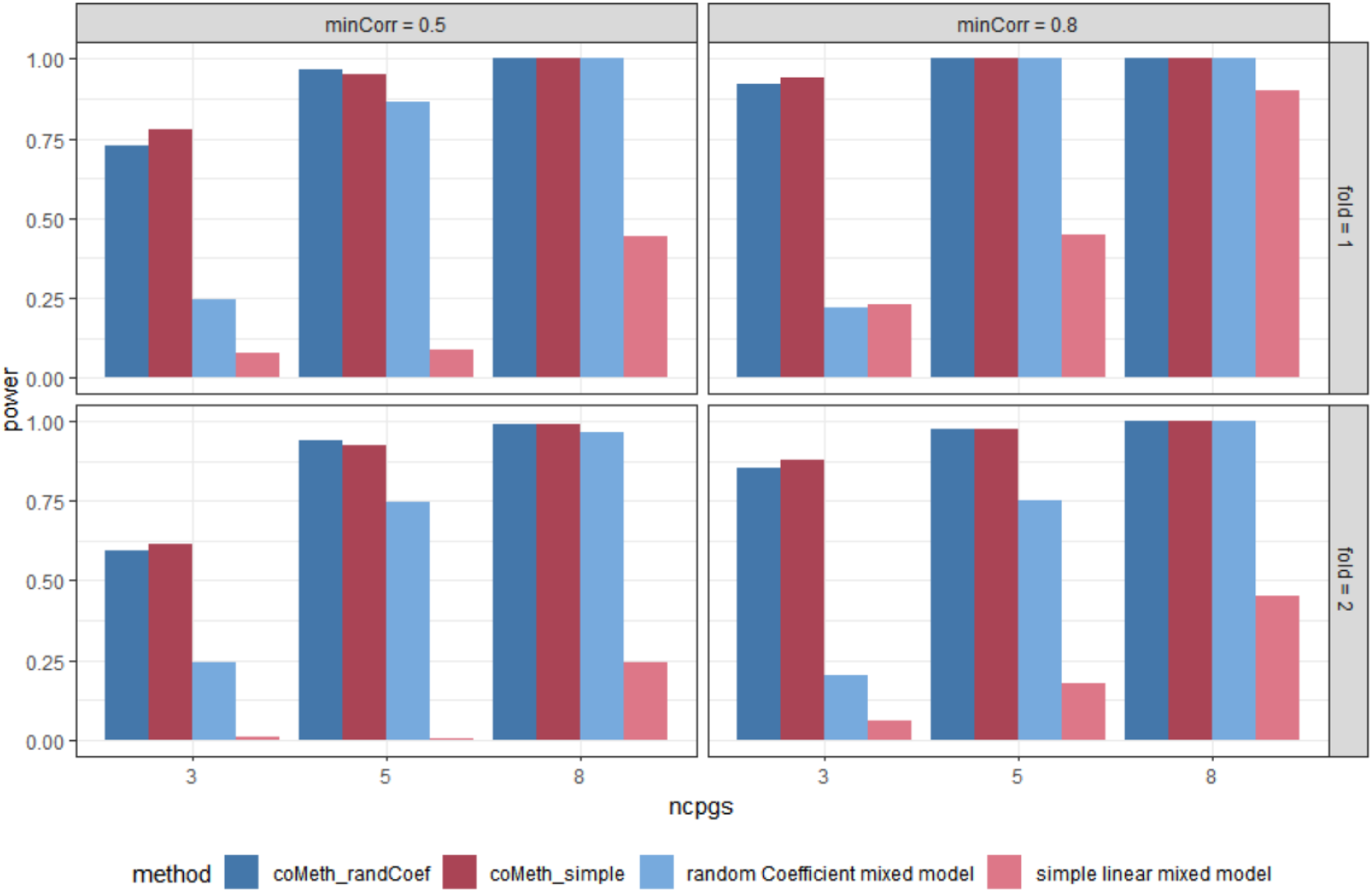
Power is improved when fitting simple linear mixed model and random coefficient mixed model to co-methylated CpGs in genomic regions (coMeth`_randCoef`, coMeth`_simple`), compared to fitting the models to all CpGs in genomic regions.

### Random coefficient mixed model improves specificity when identifying differentially methylated regions

As mentioned above, a key challenge in unsupervised approaches for identifying DMRs is their lack of specificity. In particular, it is desirable to identify significant genomic regions that include only CpG probes significantly associated with the continuous phenotype and exclude those CpGs not related to phenotype. To further evaluate the specificity of different statistical models on real methylation data, we next applied the five models described above as well as several supervised approaches (`bumphunter`(17), `DMRcate`(18), `Probe Lasso`(11), `comb-p`(10)) to all 110 prefrontal cortex samples in the Lunnon et al. dataset (7) mentioned above to identify CGIs associated with Braak scores. Braak staging scores are a standardized measure of neurofibrillary tangle burden determined at autopsy (28). These scores range from 0 to 6, indicating different pathological severity of the disease. We treat these scores as a linear variable, adjusting for covariate effects from age, sex, batch, and estimated proportions of neurons.

Supplementary Figure 2 shows mean trajectories of corrected methylation *M* values (after adjusting for covariate effects) for individual CpGs in top 10 most significant genomic regions, identified by `IMA_mean` (linear model with mean summary implemented in IMA software), `IMA_median` (linear model with median summary implemented in IMA software), `Aclust_GEE` (GEE model implemented in Aclust software), `seqlm` (simple linear mixed model implemented in `seqlm` software), `coMethDMR_simple` (simple linear mixed model implemented in `coMethDMR` software), `coMethDMR_randCoef` (random coefficient mixed model implemented in `coMethDMR` software) and comb-p software. Among the supervised methods, only comb-p returned significant regions. In the figures, each dot corresponds to an average corrected methylation *M* value for samples from a particular Braak stage. Each line represents linear regression fitted on a particular CpG. Note that within these most significant regions, there were large heterogeneities in slopes for individual CpGs for all methods, except `coMethDMR_randCoef`. Figure 8 shows standard deviations of slope estimates for individual CpGs within the top 10 regions. Each dot represents standard deviation of individual CpG slope estimates within a significant region selected by a particular method. We observed that significant regions selected by random coefficient model (`coMethDMR_randCoef`) showed much less variations in individual CpG slopes estimates by applying a linear model to single CpGs (i.e. more homogeneous associations between individual CpG methylations and disease stage).

**Figure 8.**
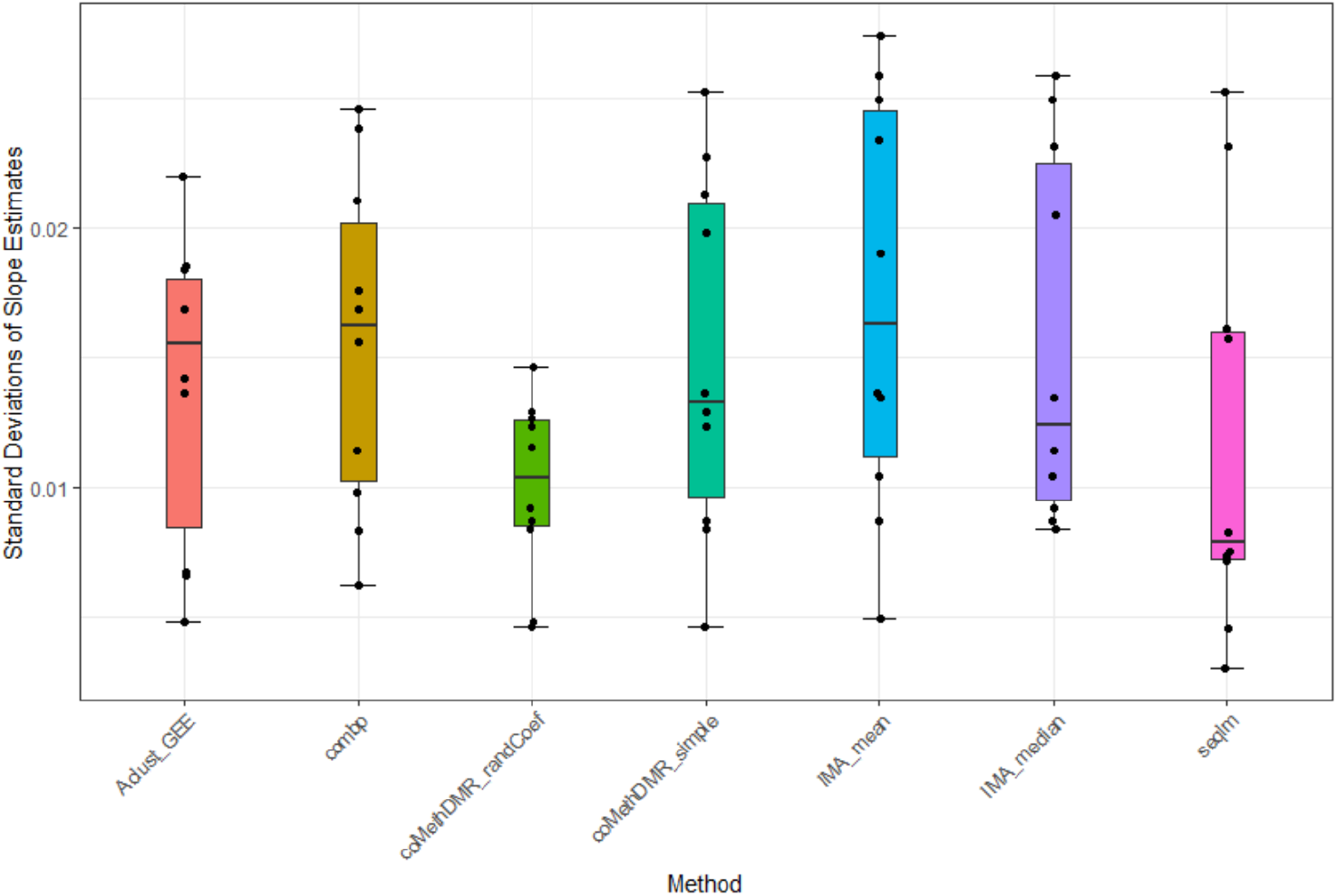
Significant regions selected by random coefficient model showed less variations in individual CpG slope estimates (i.e. more homogeneous associations between individual CpG methylations and disease stage). We considered the top 10 most significant regions by each method. Each dot represents standard deviation of individual CpG slope estimates within a significant region selected by a particular method.

To understand how the random coefficient mixed model improves specificity, note that this model specifically models co-variation of the slopes. In Figure 3 and subsection “Random coefficient mixed model”, the CpG specific slopes are modeled by random effects *b*_1*j*_, where we assume 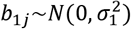. The variations in the CpG specific slopes will be captured by estimated variance component 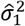 for the random effects *b*_1*j*_, which contributes to variance of 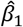, the slope main effect for the continuous phenotype (e.g. disease stage). Thus, genomic regions with more consistent differential changes in methylation levels among the CpGs will have a lower value for 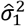, corresponding to a lower value for variance of 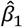, and yielding a more significant *p*-value for the slope main effect. On the other hand, regions with outlier CpGs would have large variances for 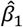, resulting non-significant *p*-values for the slope main effect.

To further illustrate the effect of modeling heterogeneity in CpG slopes, consider the 5 CpGs located within the CGI at chr13:115046754-115048034 (Figure 9). To simplify this example, we tested methylation *M* values in this region against disease stage, without controlling for any covariate variables. The results showed that the *p*-values for this region are 2.42×10^−5^ (`IMA_mean`), 0.0154 (`IMA_median`), 1.87 × 10^−4^ (GEE in Aclust), and 0.0046 (simple linear mixed model in `seqlm` and `coMethDMR_simple`). However, note that the significance of this region is driven by only 1 CpG, cg12513911 (light blue). On the other hand, the *p*-value for random coefficient model in `coMethDMR_randCoef` is 0.315, which indicates that the random coefficient mixed model correctly classified this CGI as a non-significant region.

**Figure 9.**
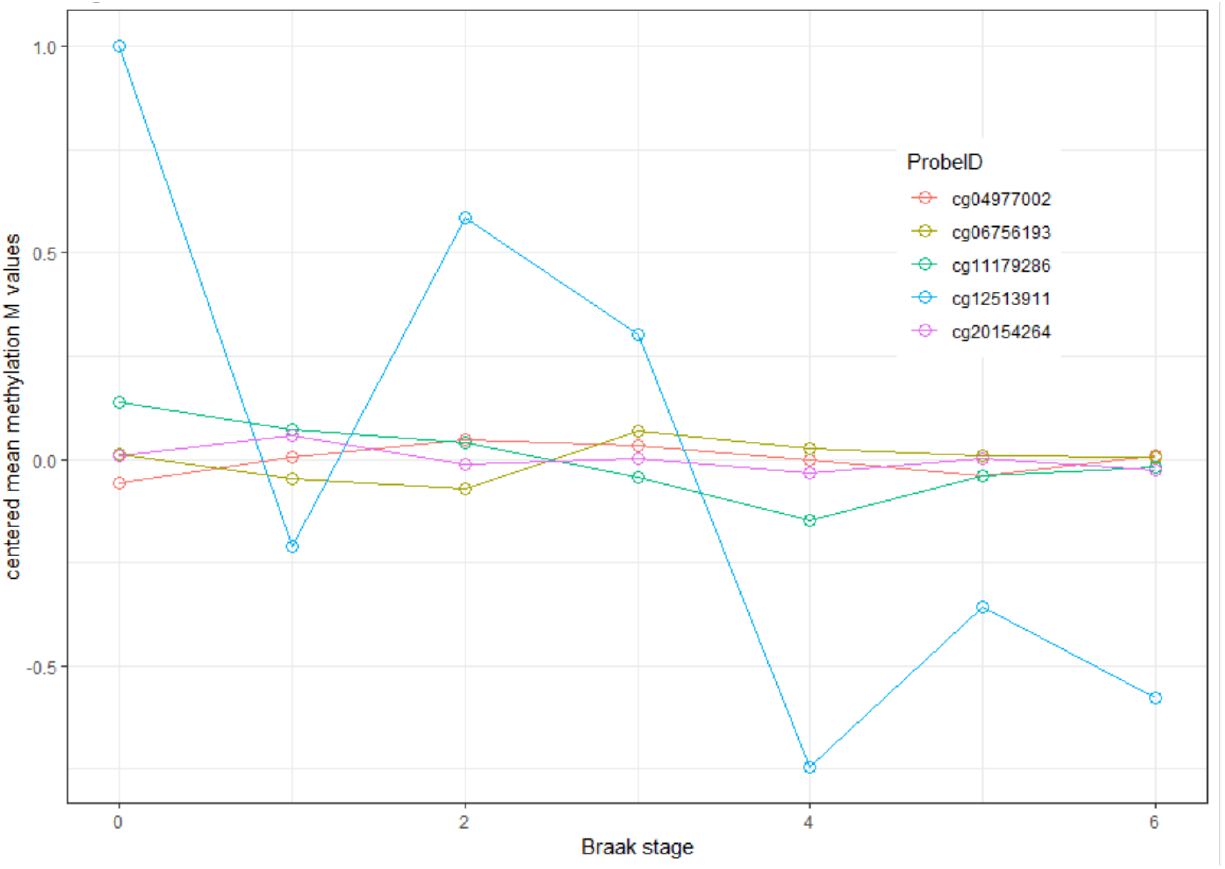
Trajectories of five individual CpGs. Each dot indicates the average methylation M value for all samples available at a given disease stage. These averages are then median centered to put all CpGs on the same graph.

This example shows that without specifically modeling variances in the slopes, the significant regions identified might have large heterogeneity in individual CpG slopes. As a result, the region may include a substantial number of non-significant CpGs. In particular, region-wise *p*-values using conventional unsupervised methods can be driven by a single outlier CpG that has strong association signal, which does not constitute a DMR by definition. In contrast, the proposed random coefficient mixed model would prioritize genomic regions where the mean methylation trajectory for multiple CpGs is highly correlated with continuous phenotype, and the heterogeneity in trajectories for individual CpGs within the region is low.

### coMethDMR identifies biologically-meaningful DMRs

To evaluate the biological plausibility of DMRs identified by different methods, we next examined the analysis results for testing methylation levels with AD stages in the Lunnon et al. dataset. After adjusting for age, sex, batch, and estimated proportions of neurons, `coMethDMR_simple` and `coMethDMR_randCoef` identified 10 and 4 significant regions at 5% false discovery rate (FDR), respectively. We compared these results with significant DMRs identified by other methods, including the `IMA_mean`, `IMA_median`, `Aclust_GEE` and `seqlm` methods. We also tested several supervised methods, including `DMRcate`, `bumphunter`, `probelasso`, and `comb-p`. Figures 10-11 show the overlap of significant regions identified by these methods and the `coMethDMR_simple` and `coMethDMR_randCoef` methods, respectively. The `seqlm` method (unsupervised) and `DMRcate`, `bumphunter`, and `probeLasso` methods (supervised) were excluded from these figures because they did not identify any DMRs at 5% FDR.

The results showed that among all other methods, results from `IMA_median` had the most overlap with both `coMethDMR_simple` and `coMethDMR_randCoef` results. In particular, half of the significant DMRs identified by `coMethDMR_simple` (5 out of 11) and `coMethDMR_randCoef` (2 out of 4) were also selected by `IMA_median` (Figure 10-11) This is most likely due to the fact that `IMA_median` is less affected by trajectories of outlying CpGs within a genomic region than other methods. Results from `Aclust_GEE` also had substantial overlap with `coMethDMR` results. However, this could be due to the fact that `Aclust_GEE` method identified a large number of DMRs. Given that the Type I error rates for `Aclust_GEE` was inflated (Figure 6), many of the significant DMRs could be false positives. Compared to the overlap between comb-p and `coMethDMR_randCoef`, results from `coMethDMR_simple` agreed more with the supervised method comb-p (Figure 12), probably because neither method accounts for heterogeneities in CpG slopes.

**Figure 10.**
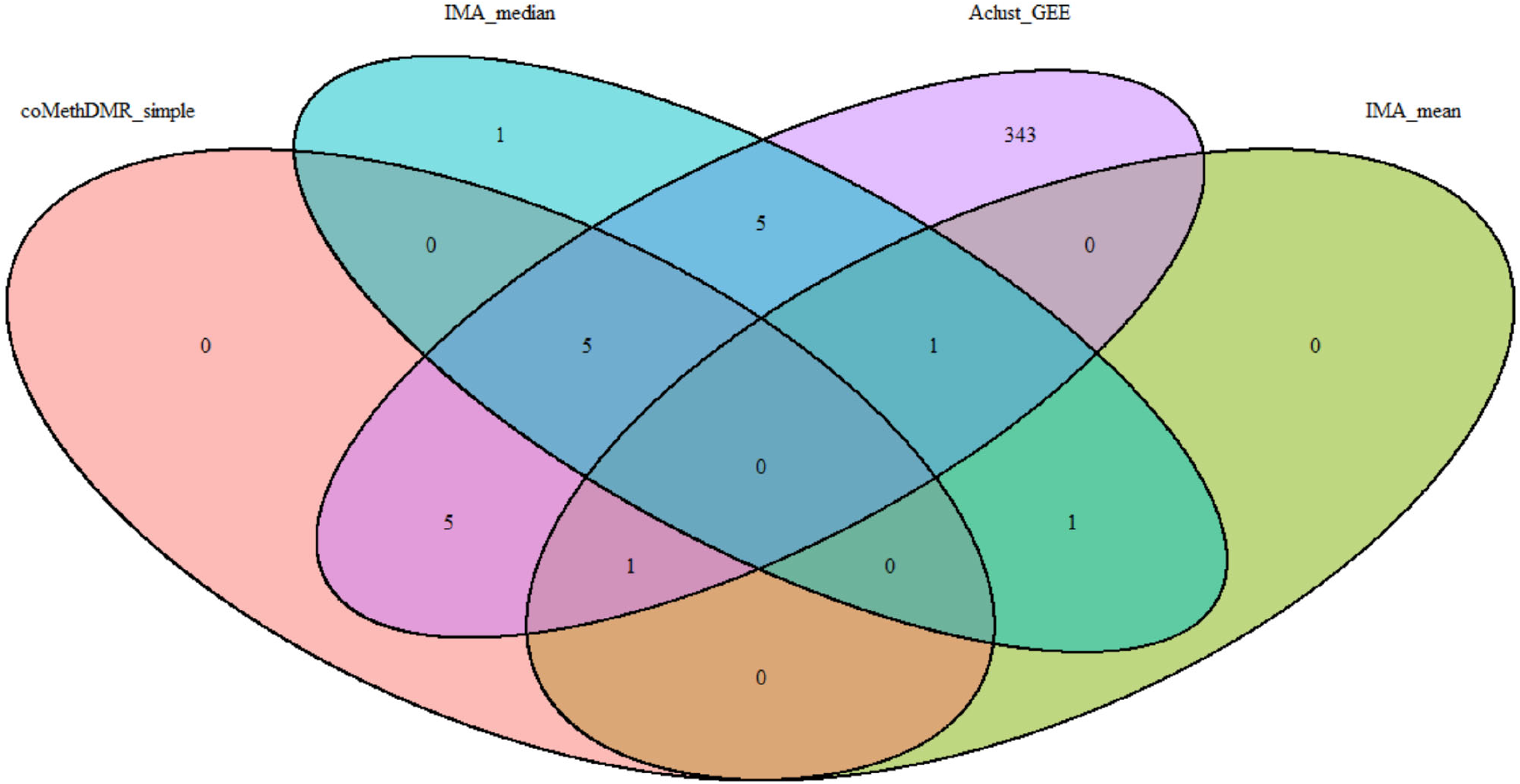
Comparison of significant regions at 5% False Discovery Rate (FDR) selected by `coMethDMR_simple` with other unsupervised approaches (`IMA_median`, `IMA_mean` and `Aclust_GEE`).

**Figure 11.**
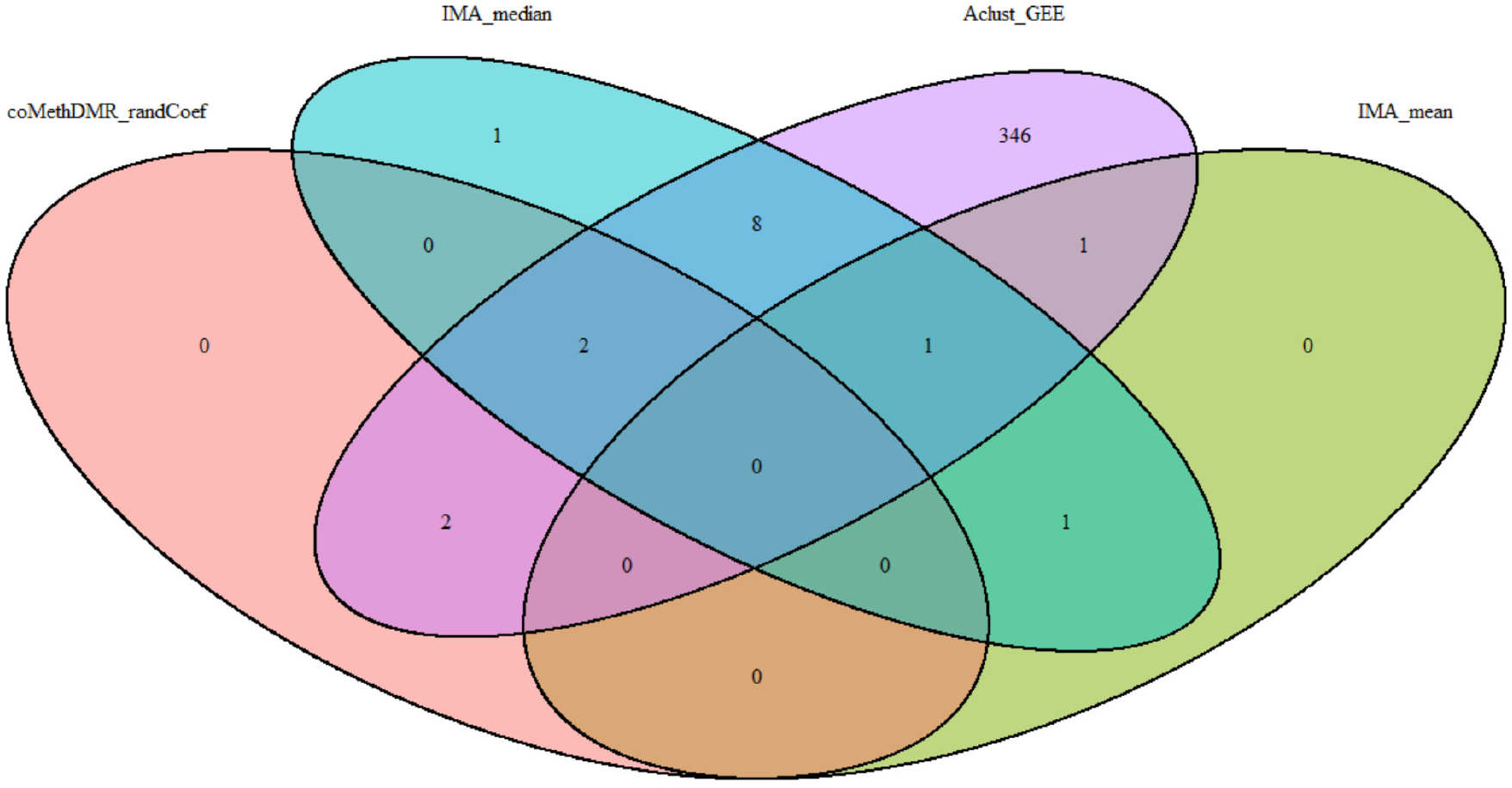
Comparison of significant regions at 5% FDR selected by `coMethDMR_randCoef` with other unsupervised approaches (`IMA_median`, `IMA_mean` and `Aclust_GEE`).

**Figure 12.**
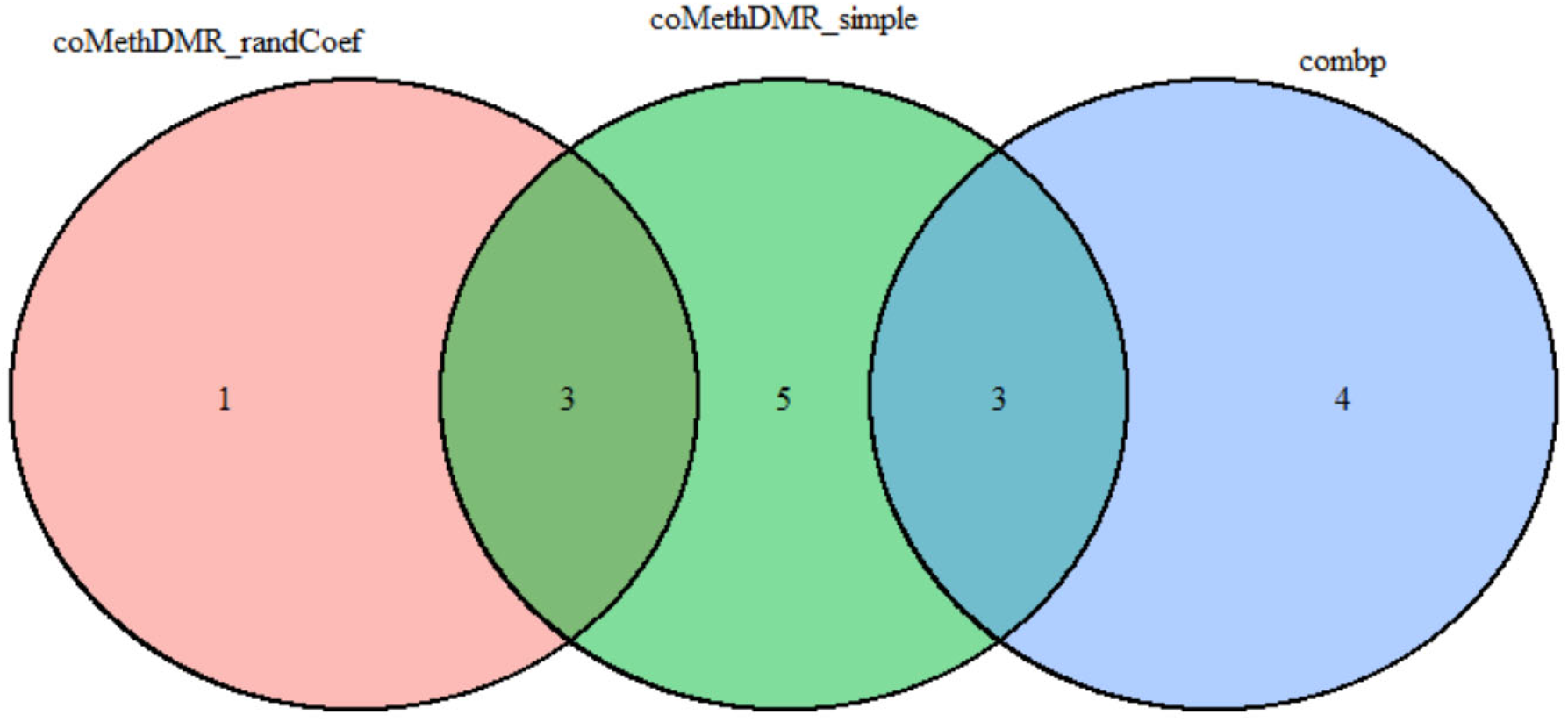
Comparison of significant regions at 5% FDR (or Sidak p-value) selected by `coMethDMR_randCoef`, `coMethDMR_simple`, and the supervised approach comb-p.

Table 2 shows the significant regions identified by `coMethDMR_simple` and `coMethDMR_randCoef`. For the DMRs identified by `coMethDMR_randCoef`, the most significant region is in the gene body of SEPT5, a brain-expressed cytoskeletal organizing gene that was nominally associated with AD (29) in family-based GWAS studies and has been shown by proteomic analysis to have altered levels in brains of AD patients (30). The second region is in KIF1A, a member of the kinesin family that transports cargo along axonal microtubules. One of KIF1A’s major roles is to transport BACE1 in neuronal axons (31). The third gene, PCNT, has also been linked to altered methylation in AD brains (6). The fourth region is in the 3’ UTR of the HOXA3 gene, which is part of a gene cluster on chromosome 7. Recently, aberrant methylation of this region has been shown to be associated with AD neuropathology in multiple AD EWAS datasets (32).

**Table 2.** Differentially methylated regions associated with AD stages identified by `coMethDMR`.

| Region | Illumina Annotation | GREAT Annotation (Distance from TSS) | Estimate | StdErr | p-Value | FDR |
| --- | --- | --- | --- | --- | --- | --- |
| **<u>method = coMethDMR randCoef</u>** |  |  |  |  |  |  |
| chr22:19709548-19709755 | SEPT5;<br>GP1BB | GP1BB (-816) | 0.050 | 0.010 | 1.38E-05 | 0.031 |
| chr2:241721922-241722113 | KIF1A | KIF1A (37707);<br>AQP12A (90756) | 0.031 | 0.007 | 1.42E-05 | 0.031 |
| chr21:47855893-47856020 | PCNT | DIP2A (-23067);<br>PCNT (111921) | 0.058 | 0.013 | 3.75E-05 | 0.044 |
| chr7:27146237-27146445 | HOXA3 | HOXA2 (0) | 0.057 | 0.013 | 4.09E-05 | 0.044 |
| **<u>method = coMethDMR simple</u>** |  |  |  |  |  |  |
| chr22:19709548-19709755 | SEPT5;<br>GP1BB | GP1BB ( -816 ) | 0.050 | 0.010 | 3.79E-06 | 0.015 |
| chr7:27153580-27153636 | HOXA3 | HOXA2 ( -11178 );<br>HOXA3 ( 5606 ) | 0.072 | 0.015 | 6.85E-06 | 0.015 |
| chr7:27146237-27146445 | HOXA3 | HOXA2 ( -3911 ) | 0.057 | 0.012 | 1.28E-05 | 0.018 |
| chr7:27185136-27185512 | HOXA6 | HOXA5 ( -2037 ) | 0.033 | 0.007 | 1.79E-05 | 0.019 |
| chr21:47855893-47856020 | PCNT | DIP2A ( -23067 );<br>PCNT ( 111921 ) | 0.058 | 0.013 | 2.52E-05 | 0.022 |
| chr17:46698881-46699073 | HOXB9 | HOXB9 ( 4862 );<br>HOXB8 ( -6676 ) | 0.042 | 0.010 | 4.23E-05 | 0.030 |
| chr18:74799495-74799572 | MBP | ZNF236 ( 263418 );<br>MBP ( 45191 ) | 0.073 | 0.017 | 4.99E-05 | 0.031 |
| chr17:74475240-74475402 | RHBDF2 | RHBDF2 ( 22168 );<br>AANAT ( 25888 ) | 0.063 | 0.015 | 6.27E-05 | 0.034 |
| chr17:3848156-3848506 | ATP2A3 | P2RX1 ( -28537 );<br>ATP2A3 ( 19254 ) | 0.049 | 0.012 | 8.21E-05 | 0.038 |
| chr7:27155002-27155358 | HOXA3 | HOXA2 ( -12750 );<br>HOXA3 ( 4034 ) | 0.048 | 0.012 | 8.85E-05 | 0.038 |
| chr16:29675846-29676071 | SPN | SPN ( 1379 );<br>QPRT ( -14399 ) | -0.055 | 0.014 | 0.000115 | 0.045 |

The `coMethDMR_simple` model identified several more genes that were previously implicated in AD etiology. For example, the MBP gene encodes myelin sheath of oligodendrocytes and Schwann cells in the nervous system. Brains from patients with AD had significant loss of intact MBP. Myelin disruption is an important feature of Alzheimer’s disease (AD) that contributes to impairment of neuronal circuitry and cognition (33,34). The RHBDF2 gene was previously found to be associated with AD stages in a different large scale methylation study (6). Another important gene, ATP2A3, encodes one of the SERCA Ca(2+)-ATPases, which are involved in maintenance of low intraneural Ca2+ concentration. The multifunction of this pump was recently found in brains of AD subjects (35). Finally, the HOXB9 and SPN genes appears to be involved in immunological and inflammatory process that may relate to AD pathophysiology (36). Taken together, these results suggest that use of `coMethDMR` can identify disease-relevant genes and replicate previous single-site and regional methylation analyses (in the case of the PCNT, RHBDF2 and HOXA3 genes).

## DISCUSSION

Although a number of methods have been proposed, identifying differentially methylated regions remains a challenging task because of the complexities in DNA methylation data. One such challenge with supervised DMR-identification methods is their lack of power when the difference in beta values between two groups was small but consistent (i.e. difference in mean beta values is less than 0.05) (25). This is likely because supervised methods scan the genome to identify regions with adjacent low p-values, so a number of positions that pass a multiple-comparison-corrected significance threshold are required. On the other hand, unsupervised methods, which define genomic regions first and then test them against phenotype, tend to lack specificity and often prioritize irrelevant genomic regions.

In this paper, we have presented `coMethDMR`, an unsupervised method for identifying differentially methylated regions for methylation data measured by Illumina arrays. Several additional features of `coMethDMR` make it especially attractive in a practical setting:

First, `coMethDMR` improves specificity and prioritizes genomic regions with co-methylated CpGs that are consistently associated with a continuous phenotype. In addition to the identification of DMRs, this improved accuracy in scoring and ranking genomic regions would also provide more accuracy in downstream analysis such as network or pathway analysis, where genes are represented by genomic regions mapped to them, as well as integration with other types of -Omics data, such as gene expression measured by RNAseq.

Second, `coMethDMR` improves power by identifying and testing co-methylated regions in the genome, instead of testing all genomic regions in the genome. By limiting analysis to only the most relevant regions in the genome, p-values are not diluted by multiple-comparison correction for regions that are unlikely to be candidate for DMRs. Note the co-methylated regions are selected without using any outcome information, so that Type I error rates for the `coMethDMR` pipeline are preserved.

Third, `coMethDMR` is flexible in the genomic regions one would like to focus on. The input for `coMethDMR` can be one, two or many genomic regions. This flexibility allows focused testing of targeted genomic regions, for example, testing significant DMRs from previous studies in a new dataset. By testing fewer number of genomic regions, the burden for multiple comparisons can be reduced. In addition to annotations provided by Illumina, other definitions of genomic regions can also be used to group CpG probes, such as the cell-type specific chromatin state segmentations identified by patterns of histone modifications in ENCODE project (39), chromatin accessible regions detected in ATAC-seq data, or transcription factor binding sites detected in ChIP-seq data. This new feature of `coMethDMR` facilitates integration of DNA methylation data with carefully-curated metadata generated by large consortia such as ENCODE (37) and Roadmap Epigenomics (38), improving power by focusing on the gene-regulatory regions which are most likely to be differentially methylated.

In summary, `coMethDMR` offers a flexible, powerful, and accurate solution for DMR analysis of array-based DNA methylation data. The entire analytical pipeline is implemented as an open-source R package, freely available to the research community. We have shown `coMethDMR` provides well-controlled false positive rate, as well as improved power over directly testing a genomic region with a continuous phenotype. In the analysis of an Alzheimer’s dataset, the agreement between results obtained by `coMethDMR` and previous reports further validates this proposed method. `coMethDMR` empowers epigenetic researchers to discover meaningful biological insights from vast amounts of large and complex DNA methylation datasets.

## AVAILABILITY

The analysis scripts used in this study, along with `coMethDMR` as an open-source R package, can be accessed at github at https://github.com/lissettegomez/coMethDMRPaper and https://github.com/lissettegomez/coMethDMR. A user guide for `coMethDMR` which provides details of commands and output is included in Supplementary Text.

## ACCESSION NUMBERS

The Lunnon et al. Alzheimer’s Disease dataset are available in the GEO data repository (accession number: GSE59685)

## ACKNOWLEDGEMENT

The authors would like to thank Michael Schmidt for testing the software and providing helpful suggestions.

---

Supplementary Table 1. Simulation study to identify optimal rdrop parameter in first step of `coMethDMR` pipeline. For each simulation scenario, rdrop parameter with red font corresponds to best performance.

Supplementary Figure 1. Type I error rates in the absence of differential methylation for different statistical models, for comethylated regions. Shown are proportions of co-methylated regions with p-values less than 0.05, for association with randomly generated “age” from Poisson distribution with mean 65, average over 10,000 simulation datasets.

Supplementary Figure 2. Mean trajectories of corrected methylation M values (after adjusting for covariate effects) for individual CpGs in top 10 most significant genomic region, identified by `IMA_mean`, `IMA_median`, `Aclust_GEE`, `seqlm`, `coMethDMR_simple`, `coMethDMR_randCoef`, and comb-p methods.

## REFERENCES

1. Portela, A. and Esteller, M. (2010) Epigenetic modifications and human disease. Nature biotechnology, 28, 1057–1068.

2. Melotte, V., Lentjes, M.H., van den Bosch, S.M., Hellebrekers, D.M., de Hoon, J.P., Wouters, K.A., Daenen, K.L., Partouns-Hendriks, I.E., Stessels, F., Louwagie, J. et al. (2009) N-Myc downstream-regulated gene 4 (NDRG4): a candidate tumor suppressor gene and potential biomarker for colorectal cancer. J Natl Cancer Inst, 101, 916–927.

3. Schmidt, B., Liebenberg, V., Dietrich, D., Schlegel, T., Kneip, C., Seegebarth, A., Flemming, N., Seemann, S., Distler, J., Lewin, J. et al. (2010) SHOX2 DNA methylation is a biomarker for the diagnosis of lung cancer based on bronchial aspirates. BMC cancer, 10, 600.

4. Jain, S., Chen, S., Chang, K.C., Lin, Y.J., Hu, C.T., Boldbaatar, B., Hamilton, J.P., Lin, S.Y., Chang, T.T., Chen, S.H. et al. (2012) Impact of the location of CpG methylation within the GSTP1 gene on its specificity as a DNA marker for hepatocellular carcinoma. PloS one, 7, e35789.

5. Lord, J. and Cruchaga, C. (2014) The epigenetic landscape of Alzheimer’s disease. Nature neuroscience, 17, 1138–1140.

6. De Jager, P.L., Srivastava, G., Lunnon, K., Burgess, J., Schalkwyk, L.C., Yu, L., Eaton, M.L., Keenan, B.T., Ernst, J., McCabe, C. et al. (2014) Alzheimer’s disease: early alterations in brain DNA methylation at ANK1, BIN1, RHBDF2 and other loci. Nature neuroscience, 17, 1156–1163.

7. Lunnon, K., Smith, R., Hannon, E., De Jager, P.L., Srivastava, G., Volta, M., Troakes, C., Al-Sarraj, S., Burrage, J., Macdonald, R. et al. (2014) Methylomic profiling implicates cortical deregulation of ANK1 in Alzheimer’s disease. Nature neuroscience, 17, 1164–1170.

8. Pidsley, R., Viana, J., Hannon, E., Spiers, H., Troakes, C., Al-Saraj, S., Mechawar, N., Turecki, G., Schalkwyk, L.C., Bray, N.J. et al. (2014) Methylomic profiling of human brain tissue supports a neurodevelopmental origin for schizophrenia. Genome biology, 15, 483.

9. Jaffe, A.E., Gao, Y., Deep-Soboslay, A., Tao, R., Hyde, T.M., Weinberger, D.R. and Kleinman, J.E. (2016) Mapping DNA methylation across development, genotype and schizophrenia in the human frontal cortex. Nature neuroscience, 19, 40–47.

10. Pedersen, B.S., Schwartz, D.A., Yang, I.V. and Kechris, K.J. (2012) Comb-p: software for combining, analyzing, grouping and correcting spatially correlated P-values. Bioinformatics, 28, 2986–2988.

11. Butcher, L.M. and Beck, S. (2015) Probe Lasso: a novel method to rope in differentially methylated regions with 450K DNA methylation data. Methods, 72, 21–28.

12. Irizarry, R.A., Ladd-Acosta, C., Carvalho, B., Wu, H., Brandenburg, S.A., Jeddeloh, J.A., Wen, B. and Feinberg, A.P. (2008) Comprehensive high-throughput arrays for relative methylation (CHARM). Genome research, 18, 780–790.

13. Jaenisch, R. and Bird, A. (2003) Epigenetic regulation of gene expression: how the genome integrates intrinsic and environmental signals. Nature genetics, 33 Suppl, 245–254.

14. Irizarry, R.A., Ladd-Acosta, C., Wen, B., Wu, Z., Montano, C., Onyango, P., Cui, H., Gabo, K., Rongione, M., Webster, M. et al. (2009) The human colon cancer methylome shows similar hypo-and hypermethylation at conserved tissue-specific CpG island shores. Nature genetics, 41, 178–186.

15. Ventham, N.T., Kennedy, N.A., Adams, A.T., Kalla, R., Heath, S., O’Leary, K.R., Drummond, H., consortium, I.B., consortium, I.C., Wilson, D.C. et al. (2016) Integrative epigenome-wide analysis demonstrates that DNA methylation may mediate genetic risk in inflammatory bowel disease. Nature communications, 7, 13507.

16. Rutten, B.P.F., Vermetten, E., Vinkers, C.H., Ursini, G., Daskalakis, N.P., Pishva, E., de Nijs, L., Houtepen, L.C., Eijssen, L., Jaffe, A.E. et al. (2018) Longitudinal analyses of the DNA methylome in deployed military servicemen identify susceptibility loci for post-traumatic stress disorder. Molecular psychiatry, 23, 1145–1156.

17. Jaffe, A.E., Murakami, P., Lee, H., Leek, J.T., Fallin, M.D., Feinberg, A.P. and Irizarry, R.A. (2012) Bump hunting to identify differentially methylated regions in epigenetic epidemiology studies. International journal of epidemiology, 41, 200–209.

18. Peters, T.J., Buckley, M.J., Statham, A.L., Pidsley, R., Samaras, K., R, V.L., Clark, S.J. and Molloy, P.L. (2015) De novo identification of differentially methylated regions in the human genome. Epigenetics & chromatin, 8, 6.

19. Kolde, R., Martens, K., Lokk, K., Laur, S. and Vilo, J. (2016) seqlm: an MDL based method for identifying differentially methylated regions in high density methylation array data. Bioinformatics, 32, 2604–2610.

20. Sofer, T., Schifano, E.D., Hoppin, J.A., Hou, L. and Baccarelli, A.A. (2013) A-clustering: a novel method for the detection of co-regulated methylation regions, and regions associated with exposure. Bioinformatics, 29, 2884–2891.

21. Chen, D.P., Lin, Y.C. and Fann, C.S. (2016) Methods for identifying differentially methylated regions for sequence-and array-based data. Briefings in functional genomics, 15, 485–490.

22. Robinson, M.D., Kahraman, A., Law, C.W., Lindsay, H., Nowicka, M., Weber, L.M. and Zhou, X. (2014) Statistical methods for detecting differentially methylated loci and regions. Frontiers in genetics, 5, 324.

23. Zhang, Q., Zhao, Y., Zhang, R., Wei, Y., Yi, H., Shao, F. and Chen, F. (2016) A Comparative Study of Five Association Tests Based on CpG Set for Epigenome-Wide Association Studies. PloS one, 11, e0156895.

24. Li, D., Xie, Z., Pape, M.L. and Dye, T. (2015) An evaluation of statistical methods for DNA methylation microarray data analysis. BMC bioinformatics, 16, 217.

25. Mallik, S., Odom, G.J., Gao, Z., Gomez, L., Chen, X. and Wang, L. (2018) An evaluation of supervised methods for identifying differentially methylated regions in Illumina methylation arrays. Briefings in bioinformatics.

26. Wang, D., Yan, L., Hu, Q., Sucheston, L.E., Higgins, M.J., Ambrosone, C.B., Johnson, C.S., Smiraglia, D.J. and Liu, S. (2012) IMA: an R package for high-throughput analysis of Illumina’s 450K Infinium methylation data. Bioinformatics, 28, 729–730.

27. Littell, R.C., Miliken, G.A., Stroup, W.W. and Wolfinger, R.D. (2006) SAS for Mixed Models. SAS Institute Inc., Cary, N.C.

28. Braak, H. and Braak, E. (1995) Staging of Alzheimer’s disease-related neurofibrillary changes. Neurobiology of aging, 16, 271–278; discussion 278-284.

29. Herold, C., Hooli, B.V., Mullin, K., Liu, T., Roehr, J.T., Mattheisen, M., Parrado, A.R., Bertram, L., Lange, C. and Tanzi, R.E. (2016) Family-based association analyses of imputed genotypes reveal genome-wide significant association of Alzheimer’s disease with OSBPL6, PTPRG, and PDCL3. Molecular psychiatry, 21, 1608–1612.

30. Musunuri, S., Wetterhall, M., Ingelsson, M., Lannfelt, L., Artemenko, K., Bergquist, J., Kultima, K. and Shevchenko, G. (2014) Quantification of the brain proteome in Alzheimer’s disease using multiplexed mass spectrometry. Journal of proteome research, 13, 2056–2068.

31. Hung, C.O. and Coleman, M.P. (2016) KIF1A mediates axonal transport of BACE1 and identification of independently moving cargoes in living SCG neurons. Traffic, 17, 1155–1167.

32. Smith, R.G., Hannon, E., De Jager, P.L., Chibnik, L., Lott, S.J., Condliffe, D., Smith, A.R., Haroutunian, V., Troakes, C., Al-Sarraj, S. et al. (2018) Elevated DNA methylation across a 48-kb region spanning the HOXA gene cluster is associated with Alzheimer’s disease neuropathology. Alzheimer’s & dementia: the journal of the Alzheimer’s Association.

33. Liao, M.C., Ahmed, M., Smith, S.O. and Van Nostrand, W.E. (2009) Degradation of amyloid beta protein by purified myelin basic protein. The Journal of biological chemistry, 284, 28917–28925.

34. Zhan, X., Jickling, G.C., Ander, B.P., Liu, D., Stamova, B., Cox, C., Jin, L.W., DeCarli, C. and Sharp, F.R. (2014) Myelin injury and degraded myelin vesicles in Alzheimer’s disease. Current Alzheimer research, 11, 232–238.

35. Mata, A.M., Berrocal, M. and Sepulveda, M.R. (2011) Impairment of the activity of the plasma membrane Ca(2)(+)-ATPase in Alzheimer’s disease. Biochemical Society transactions, 39, 819–822.

36. Hayashi, K., Ouchi, M., Endou, H. and Anzai, N. (2016) HOXB9 acts as a negative regulator of activated human T cells in response to amino acid deficiency. Immunology and cell biology, 94, 612–617.

37. Davis, C.A., Hitz, B.C., Sloan, C.A., Chan, E.T., Davidson, J.M., Gabdank, I., Hilton, J.A., Jain, K., Baymuradov, U.K., Narayanan, A.K. et al. (2018) The Encyclopedia of DNA elements (ENCODE): data portal update. Nucleic acids research, 46, D794–D801.

38. Chadwick, L.H. (2012) The NIH Roadmap Epigenomics Program data resource. Epigenomics, 4, 317–324.

