## Supplementary file for "coMethDMR: Accurate identification of co-methylated and differentially methylated regions in epigenome-wide association studies"

#### SUPPLEMENTARY METHODS

##### Unsupervised approaches for identifying DMRs

Typically, at each CpG site, methylation status is measured by the beta value, which is the ratio of the methylated signal intensity to the sum of both methylated and unmethylated signals after background subtraction. Beta values range from 0 (completely unmethylated) to 1 (fully methylated). More recent research incorporates logit-transformed beta values, known as *M* values, where $M = \log(beta/1 - beta)$. *M* values have been shown to have better statistical properties such as homoscedasticity (1) in methylation data analysis. Therefore, we use *M* values as the outcome variable in our statistical models.

Several approaches have been proposed for testing methylation levels in a predefined genomic region with phenotype ($X_i$):

(1) *Linear models*: An intuitive approach is to first summarize CpG methylation values within the set using the mean (or median) *M* value and then conduct further analysis using a linear model with these summarized methylation values (2). That is, fit the model $\bar{Y}_i = \beta_0 + \beta_1 X_i + \varepsilon_i$ to each genomic region, where $\bar{Y}_i$ is mean (or median) of methylation levels over all CpGs in the region for sample $i$. This method has been implemented in R package `IMA`.

(2) *GEE model*: Sofer et al. (2013) (3) proposed to analyze the correlated CpGs in a group using a generalized estimating equations model $Y_{ij} = \beta_0 + \beta_1 X_i + \varepsilon_{ij}$, where $Y_{ij}$ is the methylation value for CpG $j$ in sample $i$; and $\boldsymbol{\varepsilon} \sim F(0, \Sigma)$ for some mean-zero distribution *F* with covariance matrix Σ. This method has been implemented in R package `Aclust`.

(3) *Simple linear mixed model*: Kolde et al. (2016) (4) proposed to analyze the correlated CpGs in a region using a linear mixed model with a simple random intercept for each subject, that is $Y_{ij} = \beta_0 + \beta_1 X_i + U_i + \varepsilon_{ij}$; where $U_i$ is the sample random effect. This method has been implemented in R package `seqlm`.

(4) *Random coefficient mixed model*: This is the method proposed in this paper. To test for association between CpG set and stage of disease, we propose fitting a random coefficient mixed model that includes both a systematic component that models the mean for each group of CpGs, and a random component ($b_{0j}$ and $b_{1j}$) that models how each CpG varies about the group mean (Figure 3).

More specifically, the methylation levels in each genomic region with multiple CpGs is modeled by the mixed effects model

$$Y_{ij} = (\beta_0 + b_{0j}) + (\beta_1 + b_{1j}) \times Braak\ Stage_i + U_i + \varepsilon_{ij},$$

where $Y_{ij}$ is the methylation *M* value for sample $i$ at CpG $j$, and $\beta_0$ and $\beta_1$ are fixed effects that model the mean intercept and slope, respectively, for the group of CpGs.

We are interested in testing the null hypothesis $H_0\colon \beta_1 = 0$. Here, $(b_{0j}, b_{1j})^t \sim N(\mathbf{0}, \boldsymbol{G})$ is a vector of random effects that model how the intercept and slope for CpG $i$ deviate from intercept and slope of the mean trajectory. They are the *random coefficients*, where we assume an unstructured covariance matrix for the random intercept and slope deviations, allowing an arbitrary correlation between random intercept and slope. The random effects $U_1, U_2, \ldots, U_N \sim N(0, \sigma_U^2)$ model sample variations and account for correlations between CpGs within the same sample. Finally, $\varepsilon_{ij} \sim N(0, \sigma^2)$ represents variations due to measurement error.

**coMethDMR analysis of Alzheimer's Disease dataset**

Lunnon et al. (2014) (5) measured DNA methylation levels in autopsied brains of AD patients using the Infinium HumanMethylation 450K BeadChip platform. Methylation expression profiles for prefrontal cortex regions of 110 subjects from this study were downloaded from Gene Expression Omnibus database (accession number GSE59685). Normalization of methylation data was performed using the BMIQ method (6). Cell type proportions for each brain sample was estimated using the `cets` R package (7).

To assess the utility of our new method on real methylation data, we focused on testing association between methylation levels at CGIs and Braak stage, a standardized measure of neurofibrillary tangle burden determined at autopsy. The probe IDs for CpGs within each CGI, as defined by Illumnina 450K array annotation, were obtained from https://rforge.net/IMA/, under "Annotation file".

To identify contiguous co-methylated regions within each CGI, we first identified CpGs located closely on the Illumina array using the `clusterMaker` function in R package `bumphunter`, with parameter `maxGap = 200`. The 19977 predefined contiguous regions with at least 3 CpGs can be accessed from /inst/extdata folder of the `coMethDMR` package. Next, given methylation data, the `coMethAllRegions` function in `coMethDMR` was used to identify co-methylated clusters within each predefined regions, with parameter `rDrop = 0.4`.

Finally, we tested each contiguous and co-methylated region using the random coefficient mixed model (coMethDMR_randCoef), using the `lmmTestAllRegions` function. These mixed models included methylation *M* value as the outcome variable, fixed effects AD stage, covariate effects age of the patient for the brain sample, sex, methylation beadchip, estimated proportion of neurons in the sample, as well as random probe effects, and random sample effects that model correlations between multiple probes within the same sample. Similarly, we also tested each contiguous and co-methylated regions using the simple linear mixed model (coMethDMR_simple), which included all the terms in the random coefficient model except random probe effects.

**References**

1. Du, P., Zhang, X., Huang, C.C., Jafari, N., Kibbe, W.A., Hou, L. and Lin, S.M. (2010) Comparison of Beta-value and M-value methods for quantifying methylation levels by microarray analysis. *BMC bioinformatics*, **11**, 587.
2. Wang, D., Yan, L., Hu, Q., Sucheston, L.E., Higgins, M.J., Ambrosone, C.B., Johnson, C.S., Smiraglia, D.J. and Liu, S. (2012) IMA: an R package for high-throughput analysis of Illumina's 450K Infinium methylation data. *Bioinformatics*, **28**, 729-730.
3. Sofer, T., Schifano, E.D., Hoppin, J.A., Hou, L. and Baccarelli, A.A. (2013) A-clustering: a novel method for the detection of co-regulated methylation regions, and regions associated with exposure. *Bioinformatics*, **29**, 2884-2891.
4. Kolde, R., Martens, K., Lokk, K., Laur, S. and Vilo, J. (2016) seqlm: an MDL based method for identifying differentially methylated regions in high density methylation array data. *Bioinformatics*, **32**, 2604-2610.
5. Lunnon, K., Smith, R., Hannon, E., De Jager, P.L., Srivastava, G., Volta, M., Troakes, C., Al-Sarraj, S., Burrage, J., Macdonald, R. *et al.* (2014) Methylomic profiling implicates cortical deregulation of ANK1 in Alzheimer's disease. *Nature neuroscience*, **17**, 1164-1170.
6. Teschendorff, A.E., Marabita, F., Lechner, M., Bartlett, T., Tegner, J., Gomez-Cabrero, D. and Beck, S. (2013) A beta-mixture quantile normalization method for correcting probe design bias in Illumina Infinium 450 k DNA methylation data. *Bioinformatics*, **29**, 189-196.
7. Guintivano, J., Aryee, M.J. and Kaminsky, Z.A. (2013) A cell epigenotype specific model for the correction of brain cellular heterogeneity bias and its application to age, brain region and major depression. *Epigenetics*, **8**, 290-302.

### coMethDMR's User Guide

Lissette Gomez, Gabriel Odom, Lily Wang

March 11, 2019

coMethDMR is an R package that identifies genomic regions that are both co-methylated and differentially methylated in Illumina array datasets. Instead of testing all CpGs within a genomic region, coMethDMR carries out an additional step that selects co-methylated sub-regions first without using any outcome information. Next, coMethDMR tests association between methylation within the sub-region and continuous phenotype using a random coefficient mixed effects model, which models both variations between CpG sites within the region and differential methylation simultaneously. coMethDMR is available from GitHub, and will be submitted to Bioconductor soon.

#### 1. Quick start

##### 1.1 Installation

The latest version can be installed by

```
library(devtools)
install_github ("lissettegomez/coMethDMR")
```

After installation, the coMethDMR package can be loaded into R using:

```
library(coMethDMR)
```

##### 1.2 Datasets

The input of coMethDMR are methylation beta values. We assume quality control and normalization of the methylation dataset have been performed, by R packages such as minfi or RnBeads. For illustration, we use a subset of prefrontal cortex methylation data (GEO GSE59685) from a recent Alzheimer's disease epigenome-wide association study which was described in Lunnon et al. (2014). This example dataset contains beta values for 8552 CpGs on chromosome 22 for a random selection of 20 subjects.

```
data(betaMatrixChr22_df)
betaMatrixChr22_df [1:5, 1:5]

##            GSM1443279 GSM1443663 GSM1443434 GSM1443547 GSM1443577
## cg00004192  0.9249942  0.8463296  0.8700718  0.9058205  0.9090382
## cg00004775  0.6523025  0.6247554  0.7573476  0.6590817  0.6726261
## cg00012194  0.8676339  0.8679048  0.8484754  0.8754985  0.8484458
## cg00013618  0.9466056  0.9475467  0.9566493  0.9588431  0.9419563
## cg00014104  0.3932388  0.5525716  0.4075900  0.3997278  0.3216956
```

The corresponding phenotype dataset included variables `stage` (Braak AD stage), `subject.id`, `Mplate` (batch effect), `Sex`, `Sample` and `age.brain` (age of the brain donor).

```
data(pheno_df)
head(pheno_df)
```

```
##    stage subject.id     Mplate         sex     Sample age.brain
## 3      0          1 6042316048 Sex: FEMALE GSM1443251        82
## 8      2          2 6042316066 Sex: FEMALE GSM1443256        82
## 10    NA          3 6042316066   Sex: MALE GSM1443258        89
## 15     1          4 7786923107 Sex: FEMALE GSM1443263        81
## 21     2          5 6042316121 Sex: FEMALE GSM1443269        92
## 22     1          6 6042316099   Sex: MALE GSM1443270        78
```

#### 1.3 Vanilla analysis

We are interested in identifying co-methylated genomic regions associated with AD stages (`stage` treated as a linear variable). Here we illustrate analysis of genomic regions mapped to CpG islands, however the workflow can be similarly conducted for other types of genomic region as well. See section 2.1 below.

There are several steps: (1) obtain genomic regions mapped to CpG islands, (2) identify co-methylated regions, and (3) test co-methylated regions against the outcome variable AD stage.

For the first step, we use the following commands:

```
CpGisland_ls <- readRDS(
  system.file (
    "extdata",
    "CpGislandsChr22_ex.RDS",
    package = 'coMethDMR',
    mustWork = TRUE
    )
)
```

Here, `CpGisland_ls` is a list of 20 items, with each item of the list including a group of CpG probe IDs located closely within a particular CpG island region. Section 2.1 discusses how to import additional genomic regions.

Next, we identify co-methylated regions:

```
coMeth_ls <- CoMethAllRegions (
  betaMatrix = betaMatrixChr22_df,
  file = CpGisland_ls,
  fileType = "RDS",
  arrayType = "450k",
  returnAllCpGs = FALSE
)

coMeth_ls$CpGsSubregions
```

```
## $`chr22:18268062-18268249`
## [1] "cg12460175" "cg14086922" "cg21463605"
##
## $`chr22:18324579-18324769`
## [1] "cg19606103" "cg14031491" "cg03816851"
##
## $`chr22:18531243-18531447`
## [1] "cg25257671" "cg06961233" "cg08819022"
```

`coMeth_ls` is list with several components. In particular, the component `coMeth_ls$CpGsSubregions` is a list that contains groups of CpG probeIDs corresponding to co-methylated regions. Three comethylated regions were identified in this example.

If we want to look at co-methylation within the first co-methylated region:

```
WriteCorrPlot <- function (beta_mat){

  require (corrplot)
  require (coMethDMR)

  CpGs_char <- row.names (beta_mat)

  CpGsOrd_df <- OrderCpGsByLocation(
    CpGs_char, arrayType=c("450k"), output = "dataframe"
  )

  betaOrdered_mat <- t(beta_mat [CpGsOrd_df$cpg ,])

  corr <- cor (
    betaOrdered_mat, method = "spearman", use = "pairwise.complete.obs"
  )

  corrplot(corr, method="number", number.cex = 1, tl.cex = 0.7)
}

# subsetting beta values to include only co-methylated probes
betas_df <- subset(
  betaMatrixChr22_df,
  row.names(betaMatrixChr22_df) %in% coMeth_ls$CpGsSubregions[[1]]
)

WriteCorrPlot (betas_df)
```

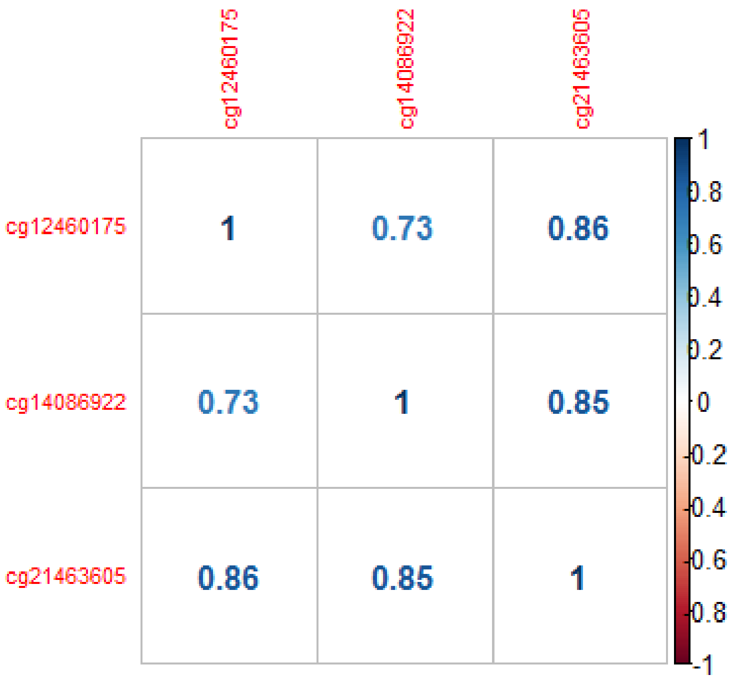

Next, we test these co-methylated regions against `stage` using a random coefficient model (more details in section 2.3 below), adjusting for `age.brain`.

```
out_df <- lmmTestAllRegions(
   beta_df = betaMatrixChr22_df,
   region_ls = coMeth_ls$CpGsSubregions,
   pheno_df,
   contPheno_char = "stage",
   covariates_char = "age.brain",
   modelType = "randCoef",
   arrayType = "450k"
 )

out_df

##   chrom    start      end nCpGs    Estimate     StdErr      Stat
## 1 chr22 18268062 18268249     3 -0.07320788 0.03943577 -1.856383
## 2 chr22 18324579 18324769     3  0.03061423 0.02953080  1.036688
## 3 chr22 18531243 18531447     3 -0.07170046 0.04631595 -1.548073
##       pValue       FDR
## 1 0.06339902 0.1824072
## 2 0.29988117 0.2998812
## 3 0.12160480 0.1824072
```

Here `out_df` is a data frame of genomic regions, with corresponding p-values and false discovery rate (FDRs) from the random coefficient mixed model.

To further examine the significant regions, we can also extract individual CpG p-values within these significant regions. For example, for the most significant region `chr22:18268062-18268249`,

```
outCpGs_df <- CpGsInfoOneRegion(
 regionName_char = "chr22:18268062-18268249",
 betas_df = betaMatrixChr22_df,
 pheno_df, contPheno_char = "stage",
 covariates_char = "age.brain",
 arrayType = "450k"
)

outCpGs_df

##                    Region        cpg   chr      pos slopeEstimate
## 1 chr22:18268062-18268249 cg12460175 chr22 18268062       -0.0387
## 2 chr22:18268062-18268249 cg14086922 chr22 18268239       -0.0795
## 3 chr22:18268062-18268249 cg21463605 chr22 18268249       -0.1015
##   slopePval UCSC_RefGene_Name UCSC_RefGene_Accession UCSC_RefGene_Group
## 1    0.3152
## 2    0.0504
## 3    0.0214
```

These CpGs mapped to intergenic regions, so there are no gene names associated with the probes. For genic regions such as `chr22:19709548-19709755`, we would have results such as the following:

```
CpGsInfoOneRegion(
  regionName_char = "chr22:19709548-19709755",
  betas_df = betaMatrixChr22_df,
  pheno_df, contPheno_char = "stage",
  covariates_char = "age.brain",
  arrayType = "450k"
)

##                    Region        cpg   chr      pos slopeEstimate
## 1 chr22:19709548-19709755 cg04533276 chr22 19709548       -0.0731
## 2 chr22:19709548-19709755 cg20193802 chr22 19709696       -0.0554
## 3 chr22:19709548-19709755 cg05726109 chr22 19709755       -0.0111
##   slopePval UCSC_RefGene_Name UCSC_RefGene_Accession UCSC_RefGene_Group
## 1    0.1229             SEPT5              NM_002688               Body
## 2    0.2097       SEPT5;GP1BB    NM_002688;NM_000407       Body;TSS1500
## 3    0.7714       SEPT5;GP1BB    NM_002688;NM_000407       Body;TSS1500
```

### 2. Details of the `coMethDMR` workflow

#### 2.1 Genomic regions

Genomic regions on the Illumina arrays can be defined based on their relations to genes or CpG Islands. Genomic regions related to genes include TSS1500, TSS200, UTR5, EXON1, GENEBODY and UTR3. Genomic regions related to CGIs are NSHORE, NSHELF, ISLAND, SSHORE and SSHELF.

In `coMethDMR` package, the probe IDs for CpGs within each genomic region, as defined by Illumnina 450K array annotation, were obtained from https://rforge.net/IMA/, under "Annotation file". From this list of pre-defined genomic regions, using the function `WriteCloseByAllRegions`, we identified clusters of CpGs located closely (i.e. the maximum separation between any two consecutive probes is 200bp; `maxGap = 200`), and we required each cluster to have at least 3 CpGs (`minCpGs = 3`).

These pre-computed genomic regions can be accessed from our GitHub repository `coMethDMRdata`. For each genomic region, a file with these clusters was created and saved with the suffix '3_200'. For example, to download genomic regions with cluster of CpGs mapped to CpG islands, we use the following commands:

```
gitHubPath_char <- "https://raw.githubusercontent.com/lissettegomez/coMethDMR
data/master/"

CpGisland_ls <- readRDS (
  url(paste0(gitHubPath_char, "ISLAND3_200.rds"))
)
```

Here `CpGisland_ls` is a list, with each item containing a character vector of CpGs IDs for a particular CpG island.

To extract clusters of close-by CpGs from pre-defined genomic regions with different values of `maxGap` and `minCpGs`, the `WriteCloseByAllRegions` function can be used.

#### 2.2 Identify co-methylated regions

Within each genomic region, we next identify contiguous and co-methylated CpGs sub-regions without using any outcome information. To select these co-methylated sub-regions, we use the `rdrop` statistic, which is the correlation between each CpG with the sum of methylation levels in all other CpGs. The default is `rDropThresh_num = 0.4`. We recommend this setting based on our simulation study. Note that higher `rDropThresh_num` values lead to fewer co-methylated regions.

For example, if we are interested in identifying co-methylated sub-region within the first genomic region in `Cgi_ls`:

```
Cgi_ls <- readRDS(
  system.file (
    "extdata",
```

```
    "CpGislandsChr22_ex.RDS",
    package = 'coMethDMR',
    mustWork = TRUE
    )
)

coMeth_ls <- CoMethAllRegions (
 betaMatrix = betaMatrixChr22_df,
 file = Cgi_ls[1],
 fileType = "RDS",
 arrayType = "450k",
 returnAllCpGs = FALSE
)

coMeth_ls

## $CpGsSubregions
## NULL
##
## $contiguousRegions
## list()
```

The results indicate there is no co-methylated sub-region within the first genomic region.

Next we look at a region (13th region in `Cgi_ls`) where there is a co-methylated sub-region:

```
coMeth_ls <- CoMethAllRegions (
 betaMatrix = betaMatrixChr22_df,
 file = Cgi_ls[13],
 fileType = "RDS",
 arrayType = "450k",
 returnAllCpGs = FALSE
)

coMeth_ls

## $CpGsSubregions
## $CpGsSubregions$`chr22:18268062-18268249`
## [1] "cg12460175" "cg14086922" "cg21463605"
##
##
## $contiguousRegions
## $contiguousRegions$`chr22:18267969-18268249`
##                    Region        CpG   Chr  MAPINFO    r_drop keep
## 1 chr22:18267969-18268249 cg18370151 chr22 18267969 0.3929400    0
## 2 chr22:18267969-18268249 cg12460175 chr22 18268062 0.8321128    1
## 3 chr22:18267969-18268249 cg14086922 chr22 18268239 0.8097174    1
## 4 chr22:18267969-18268249 cg21463605 chr22 18268249 0.8789664    1
##   keep_contiguous
## 1               0
```

```
## 2                1
## 3                1
## 4                1
```

Note that the resulting `coMeth_ls` contains two lists: the first list `coMeth_ls$CpGsSubregions` includes CpG probe IDs for each co-methylated sub-region.

The second list `coMeth_ls$contiguousRegions` shows the details on how the co-methylated region was obtained: Here `keep = 1` if `rDropThresh_num > 0.4` (i.e. a co-methylated CpG), and `keep_contigous` indicates if the probe is in a contiguous co-methylated region.

#### 2.3 Testing genomic regions against a continuous phenotype

To test association between a continuous phenotype and methylation values in a contiguous co-methylated region, two mixed models have been implemented in the function `lmmTestAllRegions`: a random coefficient mixed model (`modelType = "randCoef"`) and a simple linear mixed model (`modelType = "simple"`).

The random coefficient mixed model includes both a systematic component that models the mean for each group of CpGs, and a random component that models how each CpG varies with respect to the group mean (random probe effects). It also includes random sample effects that model correlations between multiple probes within the same sample.

More specifically, the random coefficient model is `methylation M value ~ contPheno_char + covariates_char + (1|Sample) + (contPheno_char|CpG)`. The last two terms are random intercepts and slopes for each CpG.

The simple linear mixed model includes all the terms in the random coefficient model except random probe effects.

The simple linear mixed model is

`methylation M value ~ contPheno_char + covariates_char + (1|Sample)`

To test one genomic region against the continuous phenotype `stage`, adjusting for `age.brain`:

```
lmmTestAllRegions(
  beta_df = betaMatrixChr22_df,
  region_ls = coMeth_ls$CpGsSubregions[1],
  pheno_df,
  contPheno_char = "stage",
  covariates_char = "age.brain",
  modelType = "randCoef",
  arrayType = "450k"
)

##   chrom    start      end nCpGs    Estimate     StdErr      Stat
## 1 chr22 18268062 18268249     3 -0.07320788 0.03943577 -1.856383
```

```
##       pValue         FDR
## 1 0.06339902 0.06339902
```

If we don't want to adjust for any covariate effect, we can set `covariates_char` to `NULL`:

```
lmmTestAllRegions(
  beta_df = betaMatrixChr22_df,
  region_ls = coMeth_ls$CpGsSubregions[1],
  pheno_df,
  contPheno_char = "stage",
  covariates_char = NULL,
  modelType = "randCoef",
  arrayType = "450k"
)
```

```
##   chrom    start      end nCpGs    Estimate     StdErr     Stat     pValue
## 1 chr22 18268062 18268249     3 -0.06678558 0.03883719 -1.71963 0.08549977
##          FDR
## 1 0.08549977
```

#### 3. Reference

Lunnon K, Smith R, Hannon E, De Jager PL, Srivastava G, Volta M, Troakes C, Al-Sarraj S, Burrage J, Macdonald R, et al (2014) Methylomic profiling implicates cortical deregulation of ANK1 in Alzheimer's disease. Nat Neurosci 17:1164-1170.

**Supplementary Table 1** Simulation study to identify optimal rdrop parameter in first step of coMethDMR pipeline. For each simulation scenario, rdrop parameter with red font corresponds to best performance.

| | | | fold =1 | | | | fold = 2 | | |
|---|---|---|---|---|---|---|---|---|---|
| minCorr | ncpgs | rdrop | Sensitivity | Specificity | AUC | | Sensitivity | Specificity | AUC |
| 0.5 | 3 | 0.1 | 0.968 | 0.792 | 0.890 | | 0.933 | 0.834 | 0.850 |
| 0.5 | 3 | 0.2 | 0.943 | 0.888 | 0.912 | | 0.880 | 0.924 | 0.894 |
| 0.5 | 3 | 0.3 | 0.898 | 0.947 | 0.916 | | 0.790 | 0.970 | 0.916 |
| 0.5 | 3 | 0.4 | 0.807 | 0.979 | 0.898 | | 0.644 | 0.989 | 0.907 |
| 0.5 | 3 | 0.5 | 0.638 | 0.994 | 0.855 | | 0.427 | 0.997 | 0.881 |
| 0.5 | 3 | 0.6 | 0.388 | 0.999 | 0.808 | | 0.214 | 1.000 | 0.856 |
| 0.5 | 3 | 0.7 | 0.153 | 1.000 | 0.772 | | 0.065 | 1.000 | 0.841 |
| 0.5 | 3 | 0.8 | 0.031 | 1.000 | 0.753 | | 0.012 | 1.000 | 0.835 |
| 0.5 | 3 | 0.9 | 0.002 | 1.000 | 0.750 | | 0.002 | 1.000 | 0.833 |
| 0.5 | 5 | 0.1 | 0.996 | 0.812 | 0.911 | | 0.991 | 0.844 | 0.879 |
| 0.5 | 5 | 0.2 | 0.988 | 0.919 | 0.960 | | 0.978 | 0.938 | 0.930 |
| 0.5 | 5 | 0.3 | 0.975 | 0.964 | 0.976 | | 0.959 | 0.974 | 0.960 |
| 0.5 | 5 | 0.4 | 0.956 | 0.985 | 0.961 | | 0.914 | 0.993 | 0.965 |
| 0.5 | 5 | 0.5 | 0.921 | 0.995 | 0.944 | | 0.818 | 0.997 | 0.953 |
| 0.5 | 5 | 0.6 | 0.815 | 0.999 | 0.919 | | 0.646 | 0.999 | 0.932 |
| 0.5 | 5 | 0.7 | 0.581 | 1.000 | 0.851 | | 0.363 | 1.000 | 0.881 |
| 0.5 | 5 | 0.8 | 0.219 | 1.000 | 0.777 | | 0.099 | 1.000 | 0.844 |
| 0.5 | 5 | 0.9 | 0.030 | 1.000 | 0.756 | | 0.009 | 1.000 | 0.834 |
| 0.5 | 8 | 0.1 | 1.000 | 0.843 | 0.922 | | 1.000 | 0.842 | 0.852 |
| 0.5 | 8 | 0.2 | 1.000 | 0.919 | 0.942 | | 0.997 | 0.923 | 0.928 |
| 0.5 | 8 | 0.3 | 1.000 | 0.967 | 0.963 | | 0.997 | 0.969 | 0.986 |
| 0.5 | 8 | 0.4 | 1.000 | 0.993 | 1.000 | | 0.989 | 0.992 | 0.974 |
| 0.5 | 8 | 0.5 | 0.990 | 0.999 | 1.000 | | 0.975 | 0.999 | 0.974 |
| 0.5 | 8 | 0.6 | 0.979 | 1.000 | 0.993 | | 0.937 | 1.000 | 0.971 |
| 0.5 | 8 | 0.7 | 0.891 | 1.000 | 0.968 | | 0.820 | 1.000 | 0.929 |
| 0.5 | 8 | 0.8 | 0.628 | 1.000 | 0.871 | | 0.493 | 1.000 | 0.889 |
| 0.5 | 8 | 0.9 | 0.279 | 1.000 | 0.790 | | 0.217 | 1.000 | 0.850 |
| 0.8 | 3 | 0.1 | 0.995 | 0.801 | 0.934 | | 0.982 | 0.837 | 0.878 |
| 0.8 | 3 | 0.2 | 0.988 | 0.896 | 0.972 | | 0.966 | 0.921 | 0.910 |
| 0.8 | 3 | 0.3 | 0.977 | 0.947 | 0.984 | | 0.941 | 0.968 | 0.938 |
| 0.8 | 3 | 0.4 | 0.961 | 0.976 | 0.993 | | 0.900 | 0.986 | 0.956 |
| 0.8 | 3 | 0.5 | 0.924 | 0.994 | 0.985 | | 0.823 | 0.996 | 0.952 |
| 0.8 | 3 | 0.6 | 0.862 | 0.999 | 0.954 | | 0.687 | 0.999 | 0.929 |
| 0.8 | 3 | 0.7 | 0.706 | 1.000 | 0.894 | | 0.439 | 1.000 | 0.887 |
| 0.8 | 3 | 0.8 | 0.389 | 1.000 | 0.821 | | 0.178 | 1.000 | 0.855 |
| 0.8 | 3 | 0.9 | 0.050 | 1.000 | 0.756 | | 0.027 | 1.000 | 0.838 |
| 0.8 | 5 | 0.1 | 1.000 | 0.820 | 0.900 | | 1.000 | 0.833 | 0.823 |
| 0.8 | 5 | 0.2 | 1.000 | 0.910 | 0.935 | | 1.000 | 0.910 | 1.000 |

| | | | | | | | | | |
|---|---|---|---|---|---|---|---|---|---|
| 0.8 | 5 | 0.3 | 1.000 | 0.935 | 0.935 | | 0.975 | 0.980 | 1.000 |
| 0.8 | 5 | 0.4 | 1.000 | 0.980 | 1.000 | | 0.955 | 0.985 | 0.976 |
| 0.8 | 5 | 0.5 | 1.000 | 0.985 | 1.000 | | 0.905 | 0.995 | 0.944 |
| 0.8 | 5 | 0.6 | 1.000 | 0.995 | 1.000 | | 0.870 | 1.000 | 0.944 |
| 0.8 | 5 | 0.7 | 0.990 | 0.995 | 1.000 | | 0.765 | 1.000 | 0.900 |
| 0.8 | 5 | 0.8 | 0.885 | 1.000 | 0.955 | | 0.560 | 1.000 | 0.900 |
| 0.8 | 5 | 0.9 | 0.245 | 1.000 | 0.000 | | 0.115 | 1.000 | 0.000 |
| 0.8 | 8 | 0.1 | 1.000 | 0.813 | 0.896 | | 1.000 | 0.834 | 0.975 |
| 0.8 | 8 | 0.2 | 1.000 | 0.938 | 0.952 | | 1.000 | 0.953 | 0.975 |
| 0.8 | 8 | 0.3 | 1.000 | 0.975 | 0.975 | | 1.000 | 0.981 | 0.975 |
| 0.8 | 8 | 0.4 | 1.000 | 0.994 | 1.000 | | 1.000 | 0.991 | 0.975 |
| 0.8 | 8 | 0.5 | 1.000 | 1.000 | 1.000 | | 1.000 | 0.997 | 1.000 |
| 0.8 | 8 | 0.6 | 1.000 | 1.000 | 1.000 | | 1.000 | 1.000 | 1.000 |
| 0.8 | 8 | 0.7 | 1.000 | 1.000 | 1.000 | | 0.961 | 1.000 | 0.985 |
| 0.8 | 8 | 0.8 | 0.989 | 1.000 | 1.000 | | 0.878 | 1.000 | 0.890 |
| 0.8 | 8 | 0.9 | 0.701 | 1.000 | 0.881 | | 0.583 | 1.000 | 0.890 |

**Suppl Fig 1.** Type I error rates in the absence of differential methylation for different statistical models, for co-methylated regions. Shown are proportions of co-methylated regions with p-values less than 0.05, for association with randomly generated “age” from Poisson distribution with mean 65, average over 10,000 simulation datasets.

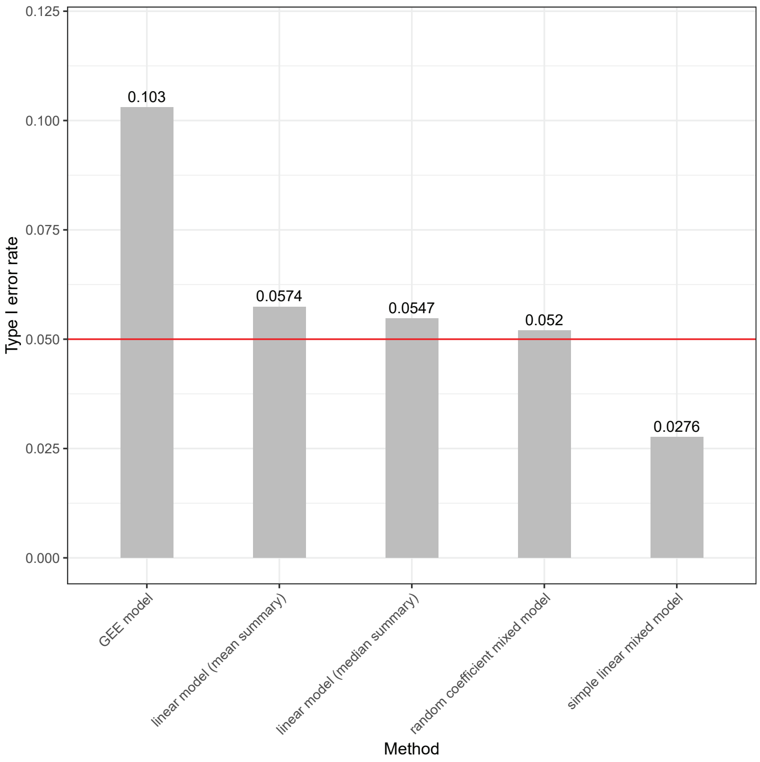

**Suppl Fig 2:** mean trajectories of corrected methylation M values (after adjusting for covariate effects) for individual CpGs in top 10 most significant genomic region, identified by IMA_mean, IMA_median, Aclust_GEE, seqlm, coMethDMR_simple, coMethDMR_randCoef and comb-p methods.

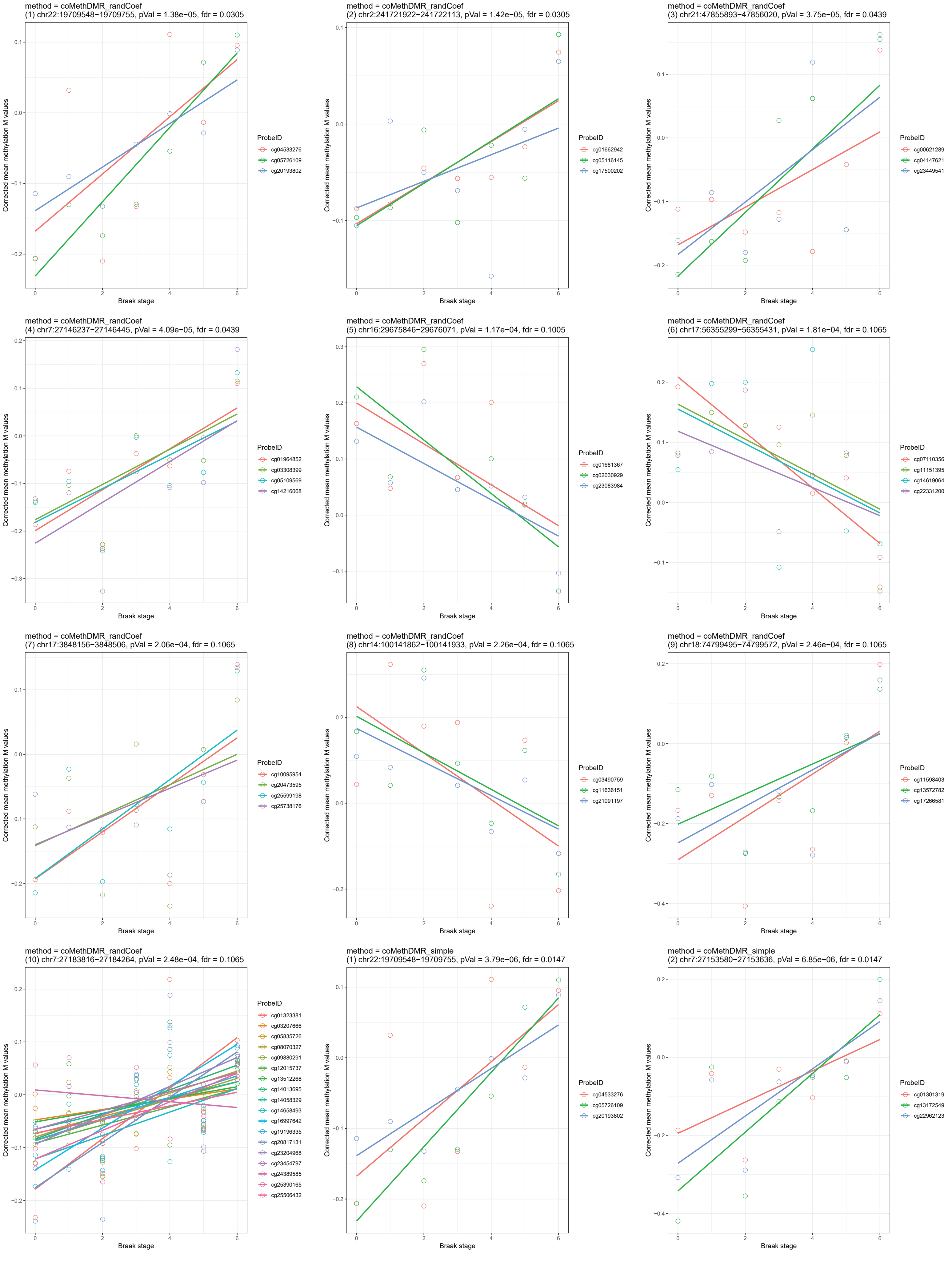

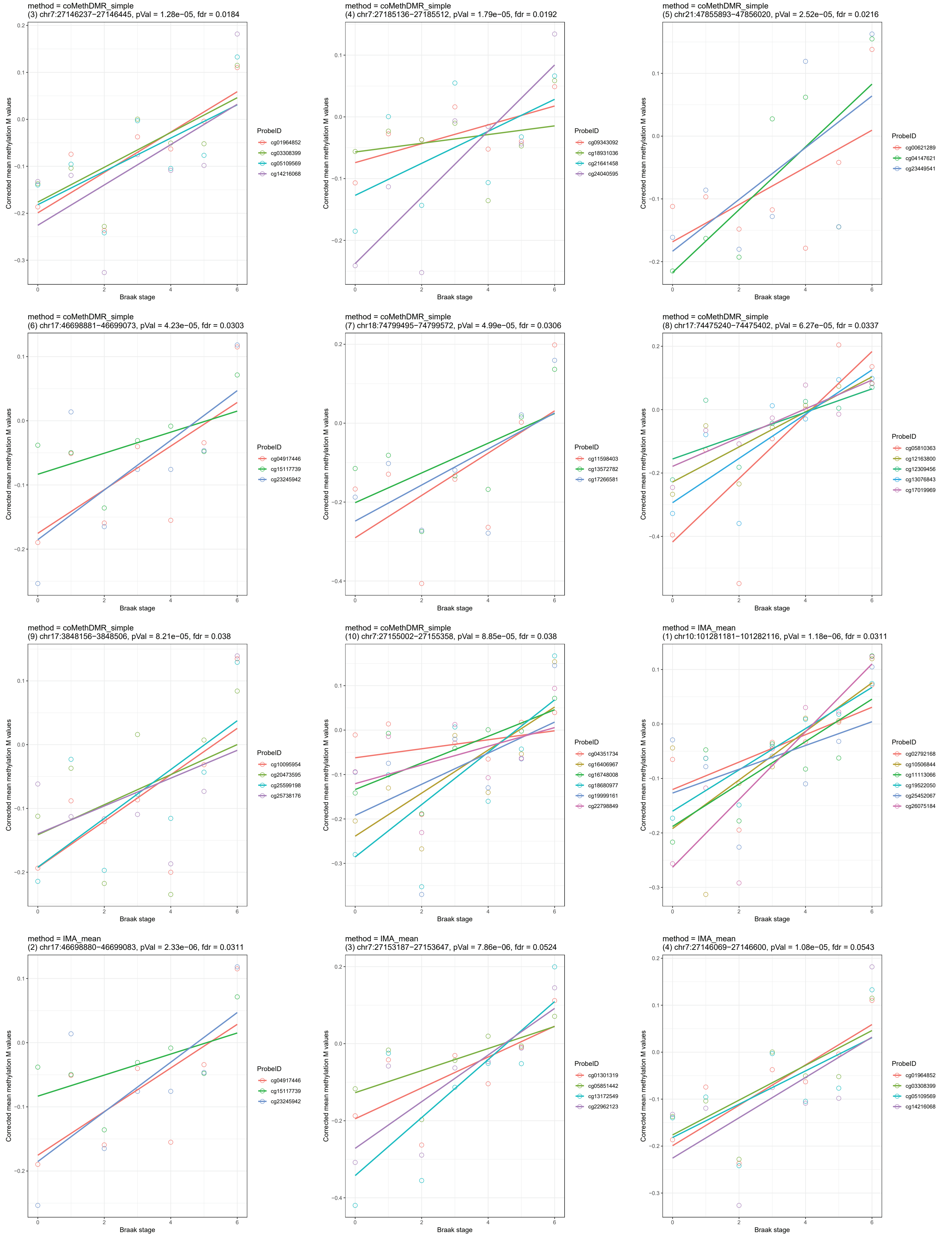
method = coMethDMR_simple
(3) chr7:27146237−27146445, pVal = 1.28e−05, fdr = 0.0184
ProbeID
cg01964852
cg03308399
cg05109569
cg14216068
method = coMethDMR_simple
(4) chr7:27185136−27185512, pVal = 1.79e−05, fdr = 0.0192
ProbeID
cg09343092
cg18931036
cg21641458
cg24040595
method = coMethDMR_simple
(5) chr21:47855893−47856020, pVal = 2.52e−05, fdr = 0.0216
ProbeID
cg00621289
cg04147621
cg23449541
method = coMethDMR_simple
(6) chr17:46698881−46699073, pVal = 4.23e−05, fdr = 0.0303
ProbeID
cg04917446
cg15117739
cg23245942
method = coMethDMR_simple
(7) chr18:74799495−74799572, pVal = 4.99e−05, fdr = 0.0306
ProbeID
cg11598403
cg13572782
cg17266581
method = coMethDMR_simple
(8) chr17:74475240−74475402, pVal = 6.27e−05, fdr = 0.0337
ProbeID
cg05810363
cg12163800
cg12309456
cg13076843
cg17019969
method = coMethDMR_simple
(9) chr17:3848156−3848506, pVal = 8.21e−05, fdr = 0.038
ProbeID
cg10095954
cg20473595
cg25599198
cg25738176
method = coMethDMR_simple
(10) chr7:27155002−27155358, pVal = 8.85e−05, fdr = 0.038
ProbeID
cg04351734
cg16406967
cg16748008
cg18680977
cg19999161
cg22798849
method = IMA_mean
(1) chr10:101281181−101282116, pVal = 1.18e−06, fdr = 0.0311
ProbeID
cg02792168
cg10506844
cg11113066
cg19522050
cg25452067
cg26075184
method = IMA_mean
(2) chr17:46698880−46699083, pVal = 2.33e−06, fdr = 0.0311
ProbeID
cg04917446
cg15117739
cg23245942
method = IMA_mean
(3) chr7:27153187−27153647, pVal = 7.86e−06, fdr = 0.0524
ProbeID
cg01301319
cg05851442
cg13172549
cg22962123
method = IMA_mean
(4) chr7:27146069−27146600, pVal = 1.08e−05, fdr = 0.0543
ProbeID
cg01964852
cg03308399
cg05109569
cg14216068
Corrected mean methylation M values
Braak stage

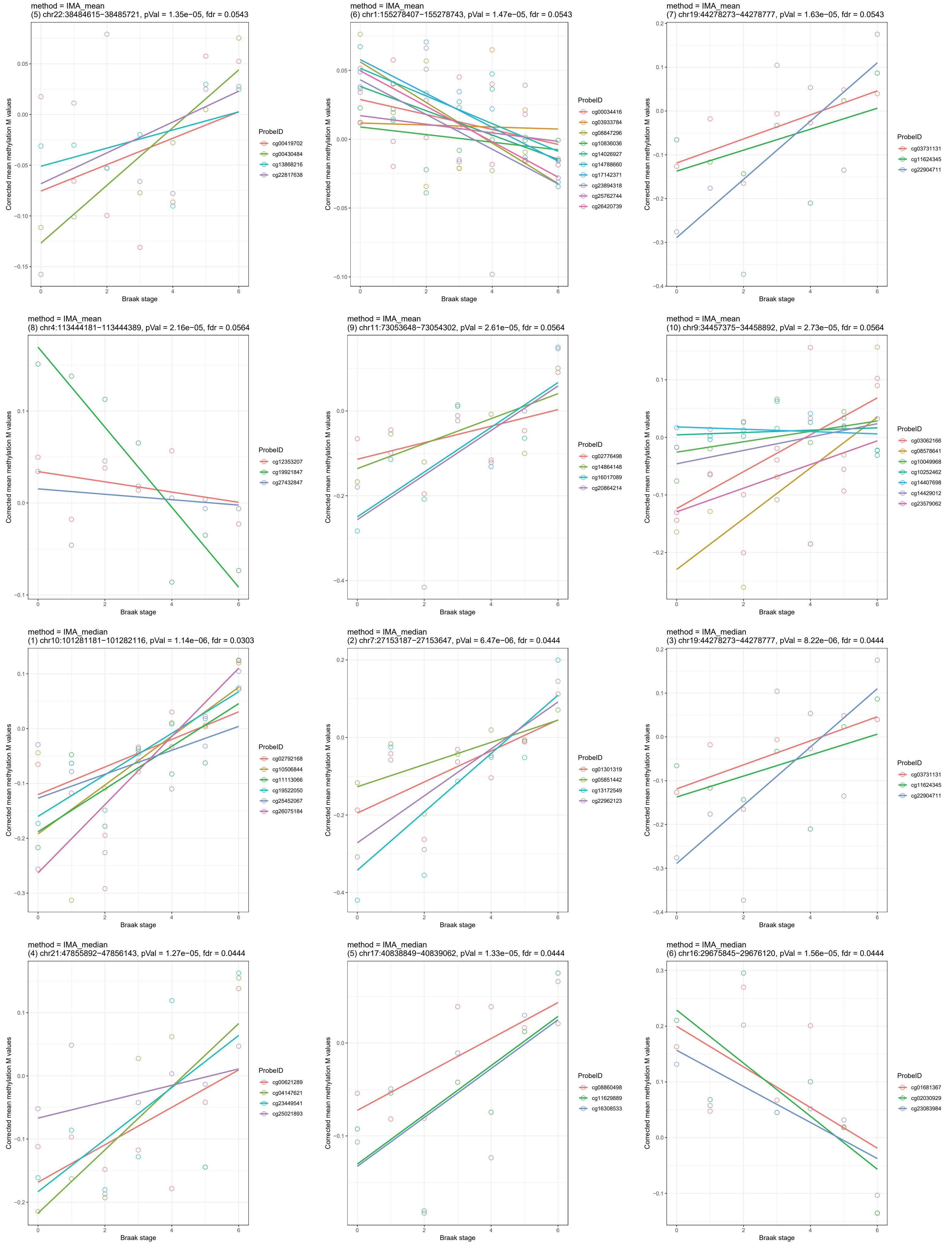

method = IMA_mean
(5) chr22:38484615−38485721, pVal = 1.35e−05, fdr = 0.0543
Corrected mean methylation M values
Braak stage
ProbeID
cg00419702
cg00430484
cg13868216
cg22817638
method = IMA_mean
(6) chr1:155278407−155278743, pVal = 1.47e−05, fdr = 0.0543
Corrected mean methylation M values
Braak stage
ProbeID
cg00034416
cg03933784
cg08847296
cg10836036
cg14026927
cg14788660
cg17142371
cg23894318
cg25762744
cg26420739
method = IMA_mean
(7) chr19:44278273−44278777, pVal = 1.63e−05, fdr = 0.0543
Corrected mean methylation M values
Braak stage
ProbeID
cg03731131
cg11624345
cg22904711
method = IMA_mean
(8) chr4:113444181−113444389, pVal = 2.16e−05, fdr = 0.0564
Corrected mean methylation M values
Braak stage
ProbeID
cg12353207
cg19921847
cg27432847
method = IMA_mean
(9) chr11:73053648−73054302, pVal = 2.61e−05, fdr = 0.0564
Corrected mean methylation M values
Braak stage
ProbeID
cg02776498
cg14864148
cg16017089
cg20864214
method = IMA_mean
(10) chr9:34457375−34458892, pVal = 2.73e−05, fdr = 0.0564
Corrected mean methylation M values
Braak stage
ProbeID
cg03062166
cg08578641
cg10049968
cg10252462
cg14407698
cg14429012
cg23579062
method = IMA_median
(1) chr10:101281181−101282116, pVal = 1.14e−06, fdr = 0.0303
Corrected mean methylation M values
Braak stage
ProbeID
cg02792168
cg10506844
cg11113066
cg19522050
cg25452067
cg26075184
method = IMA_median
(2) chr7:27153187−27153647, pVal = 6.47e−06, fdr = 0.0444
Corrected mean methylation M values
Braak stage
ProbeID
cg01301319
cg05851442
cg13172549
cg22962123
method = IMA_median
(3) chr19:44278273−44278777, pVal = 8.22e−06, fdr = 0.0444
Corrected mean methylation M values
Braak stage
ProbeID
cg03731131
cg11624345
cg22904711
method = IMA_median
(4) chr21:47855892−47856143, pVal = 1.27e−05, fdr = 0.0444
Corrected mean methylation M values
Braak stage
ProbeID
cg00621289
cg04147621
cg23449541
cg25021893
method = IMA_median
(5) chr17:40838849−40839062, pVal = 1.33e−05, fdr = 0.0444
Corrected mean methylation M values
Braak stage
ProbeID
cg08860498
cg11629889
cg16308533
method = IMA_median
(6) chr16:29675845−29676120, pVal = 1.56e−05, fdr = 0.0444
Corrected mean methylation M values
Braak stage
ProbeID
cg01681367
cg02030929
cg23083984

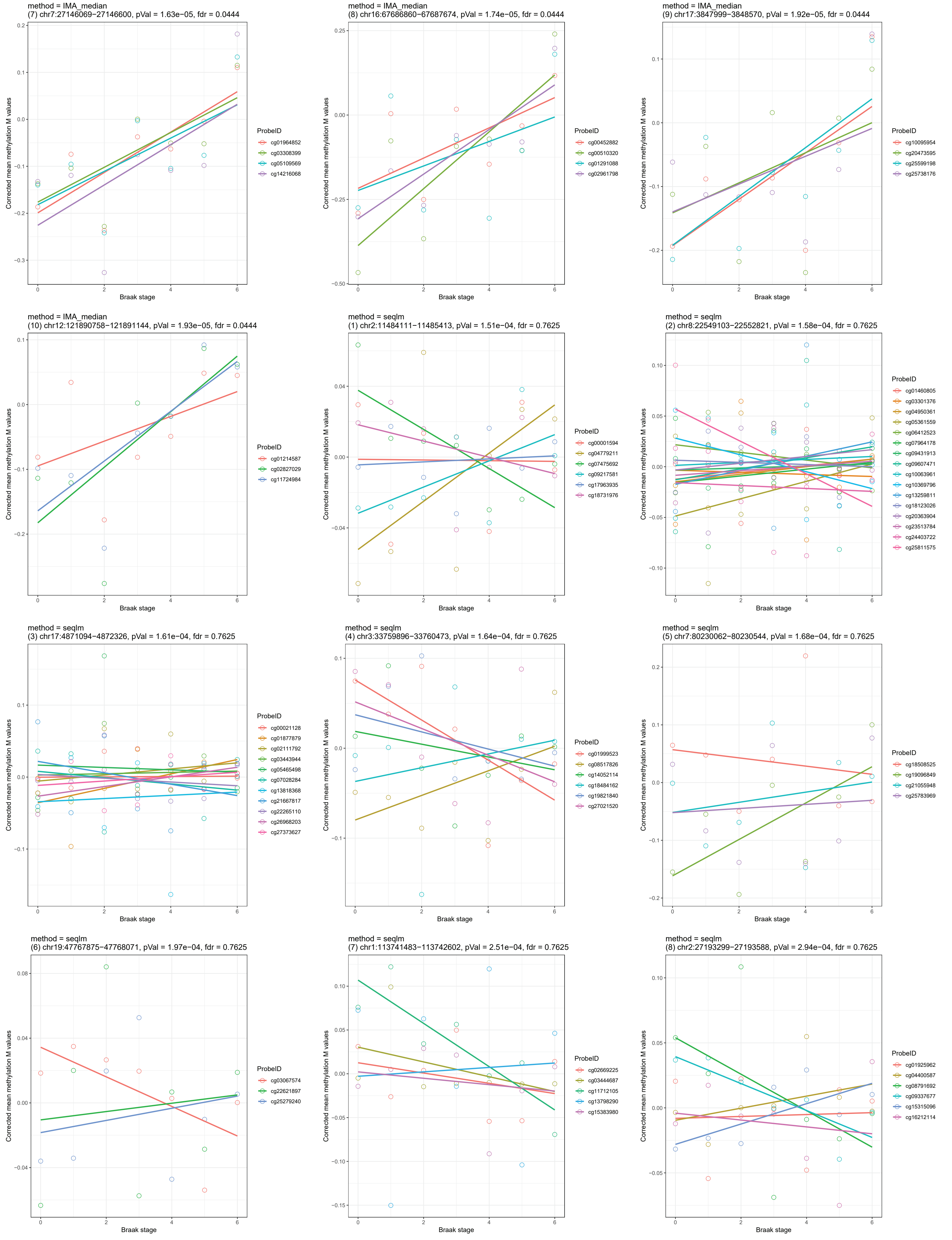

method = IMA_median
(7) chr7:27146069−27146600, pVal = 1.63e−05, fdr = 0.0444
Corrected mean methylation M values
Braak stage
ProbeID
cg01964852
cg03308399
cg05109569
cg14216068
method = IMA_median
(8) chr16:67686860−67687674, pVal = 1.74e−05, fdr = 0.0444
Corrected mean methylation M values
Braak stage
ProbeID
cg00452882
cg00510320
cg01291088
cg02961798
method = IMA_median
(9) chr17:3847999−3848570, pVal = 1.92e−05, fdr = 0.0444
Corrected mean methylation M values
Braak stage
ProbeID
cg10095954
cg20473595
cg25599198
cg25738176
method = IMA_median
(10) chr12:121890758−121891144, pVal = 1.93e−05, fdr = 0.0444
Corrected mean methylation M values
Braak stage
ProbeID
cg01214587
cg02827029
cg11724984
method = seqlm
(1) chr2:11484111−11485413, pVal = 1.51e−04, fdr = 0.7625
Corrected mean methylation M values
Braak stage
ProbeID
cg00001594
cg04779211
cg07475692
cg09217581
cg17963935
cg18731976
method = seqlm
(2) chr8:22549103−22552821, pVal = 1.58e−04, fdr = 0.7625
Corrected mean methylation M values
Braak stage
ProbeID
cg01460805
cg03301376
cg04950361
cg05361559
cg06412523
cg07964178
cg09431913
cg09607471
cg10063961
cg10369796
cg13259811
cg18123026
cg20363904
cg23513784
cg24403722
cg25811575
method = seqlm
(3) chr17:4871094−4872326, pVal = 1.61e−04, fdr = 0.7625
Corrected mean methylation M values
Braak stage
ProbeID
cg00021128
cg01877879
cg02111792
cg03443944
cg05465498
cg07028284
cg13818368
cg21667817
cg22265110
cg26968203
cg27373627
method = seqlm
(4) chr3:33759896−33760473, pVal = 1.64e−04, fdr = 0.7625
Corrected mean methylation M values
Braak stage
ProbeID
cg01999523
cg08517826
cg14052114
cg18484162
cg19821840
cg27021520
method = seqlm
(5) chr7:80230062−80230544, pVal = 1.68e−04, fdr = 0.7625
Corrected mean methylation M values
Braak stage
ProbeID
cg18508525
cg19096849
cg21055948
cg25783969
method = seqlm
(6) chr19:47767875−47768071, pVal = 1.97e−04, fdr = 0.7625
Corrected mean methylation M values
Braak stage
ProbeID
cg03067574
cg22621897
cg25279240
method = seqlm
(7) chr1:113741483−113742602, pVal = 2.51e−04, fdr = 0.7625
Corrected mean methylation M values
Braak stage
ProbeID
cg02669225
cg03444687
cg11712105
cg13798290
cg15383980
method = seqlm
(8) chr2:27193299−27193588, pVal = 2.94e−04, fdr = 0.7625
Corrected mean methylation M values
Braak stage
ProbeID
cg01925962
cg04400587
cg08791692
cg09337677
cg15315096
cg16212114

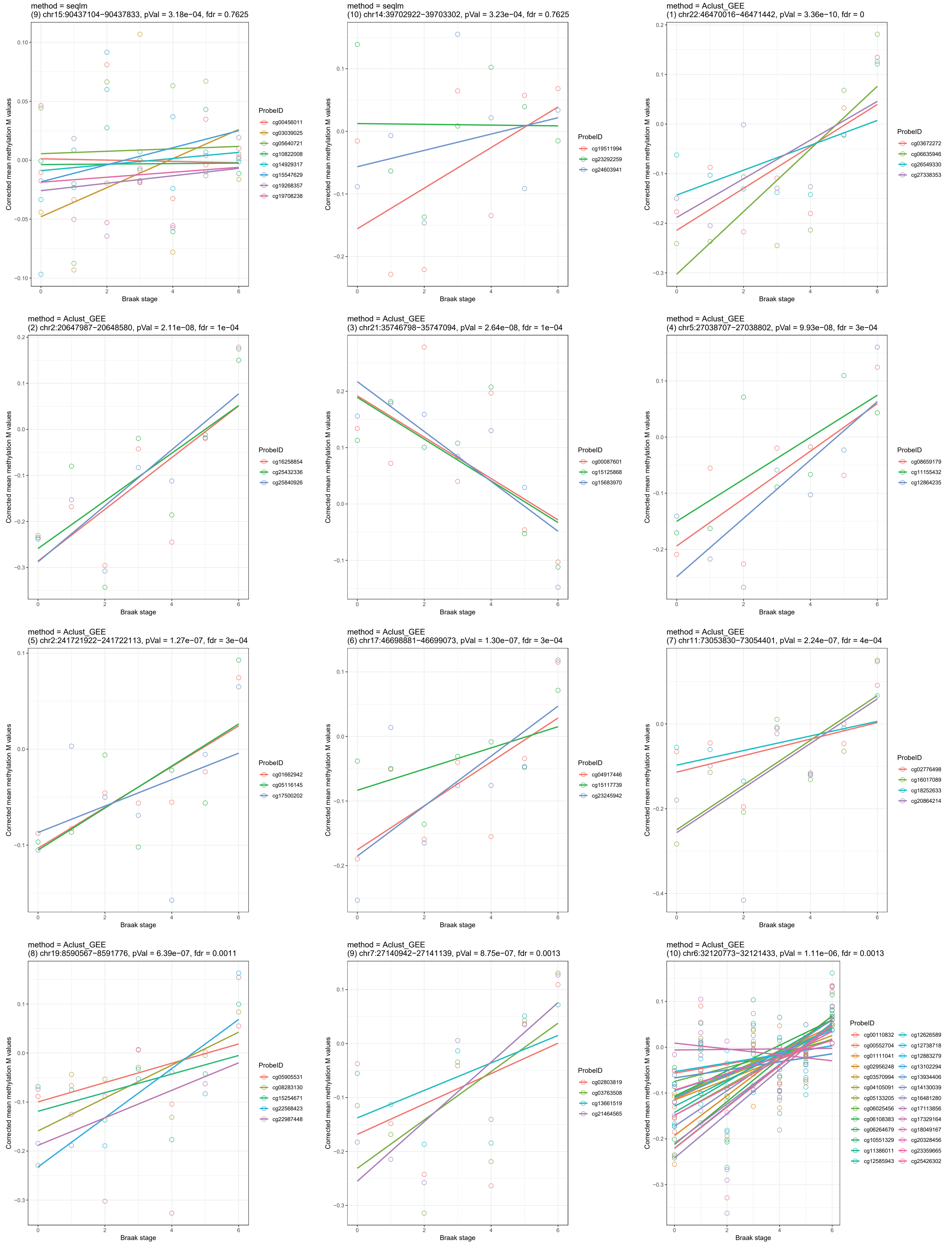
method = seqlm
(9) chr15:90437104−90437833, pVal = 3.18e−04, fdr = 0.7625
Corrected mean methylation M values
Braak stage
ProbeID
cg00456011
cg03039025
cg05640721
cg10822008
cg14929317
cg15547629
cg19268357
cg19708238
method = seqlm
(10) chr14:39702922−39703302, pVal = 3.23e−04, fdr = 0.7625
Corrected mean methylation M values
Braak stage
ProbeID
cg19511994
cg23292259
cg24603941
method = Aclust_GEE
(1) chr22:46470016−46471442, pVal = 3.36e−10, fdr = 0
Corrected mean methylation M values
Braak stage
ProbeID
cg03672272
cg06635946
cg26549330
cg27338353
method = Aclust_GEE
(2) chr2:20647987−20648580, pVal = 2.11e−08, fdr = 1e−04
Corrected mean methylation M values
Braak stage
ProbeID
cg16258854
cg25432336
cg25840926
method = Aclust_GEE
(3) chr21:35746798−35747094, pVal = 2.64e−08, fdr = 1e−04
Corrected mean methylation M values
Braak stage
ProbeID
cg00087601
cg15125868
cg15683970
method = Aclust_GEE
(4) chr5:27038707−27038802, pVal = 9.93e−08, fdr = 3e−04
Corrected mean methylation M values
Braak stage
ProbeID
cg08659179
cg11155432
cg12864235
method = Aclust_GEE
(5) chr2:241721922−241722113, pVal = 1.27e−07, fdr = 3e−04
Corrected mean methylation M values
Braak stage
ProbeID
cg01662942
cg05116145
cg17500202
method = Aclust_GEE
(6) chr17:46698881−46699073, pVal = 1.30e−07, fdr = 3e−04
Corrected mean methylation M values
Braak stage
ProbeID
cg04917446
cg15117739
cg23245942
method = Aclust_GEE
(7) chr11:73053830−73054401, pVal = 2.24e−07, fdr = 4e−04
Corrected mean methylation M values
Braak stage
ProbeID
cg02776498
cg16017089
cg18252633
cg20864214
method = Aclust_GEE
(8) chr19:8590567−8591776, pVal = 6.39e−07, fdr = 0.0011
Corrected mean methylation M values
Braak stage
ProbeID
cg05905531
cg08283130
cg15254671
cg22568423
cg22987448
method = Aclust_GEE
(9) chr7:27140942−27141139, pVal = 8.75e−07, fdr = 0.0013
Corrected mean methylation M values
Braak stage
ProbeID
cg02803819
cg03763508
cg13661519
cg21464565
method = Aclust_GEE
(10) chr6:32120773−32121433, pVal = 1.11e−06, fdr = 0.0013
Corrected mean methylation M values
Braak stage
ProbeID
cg00110832
cg00552704
cg01111041
cg02956248
cg03570994
cg04105091
cg05133205
cg06025456
cg06108383
cg06264679
cg10551329
cg11386011
cg12585943
cg12626589
cg12738718
cg12883279
cg13102294
cg13934406
cg14130039
cg16481280
cg17113856
cg17329164
cg18049167
cg20328456
cg23359665
cg25426302

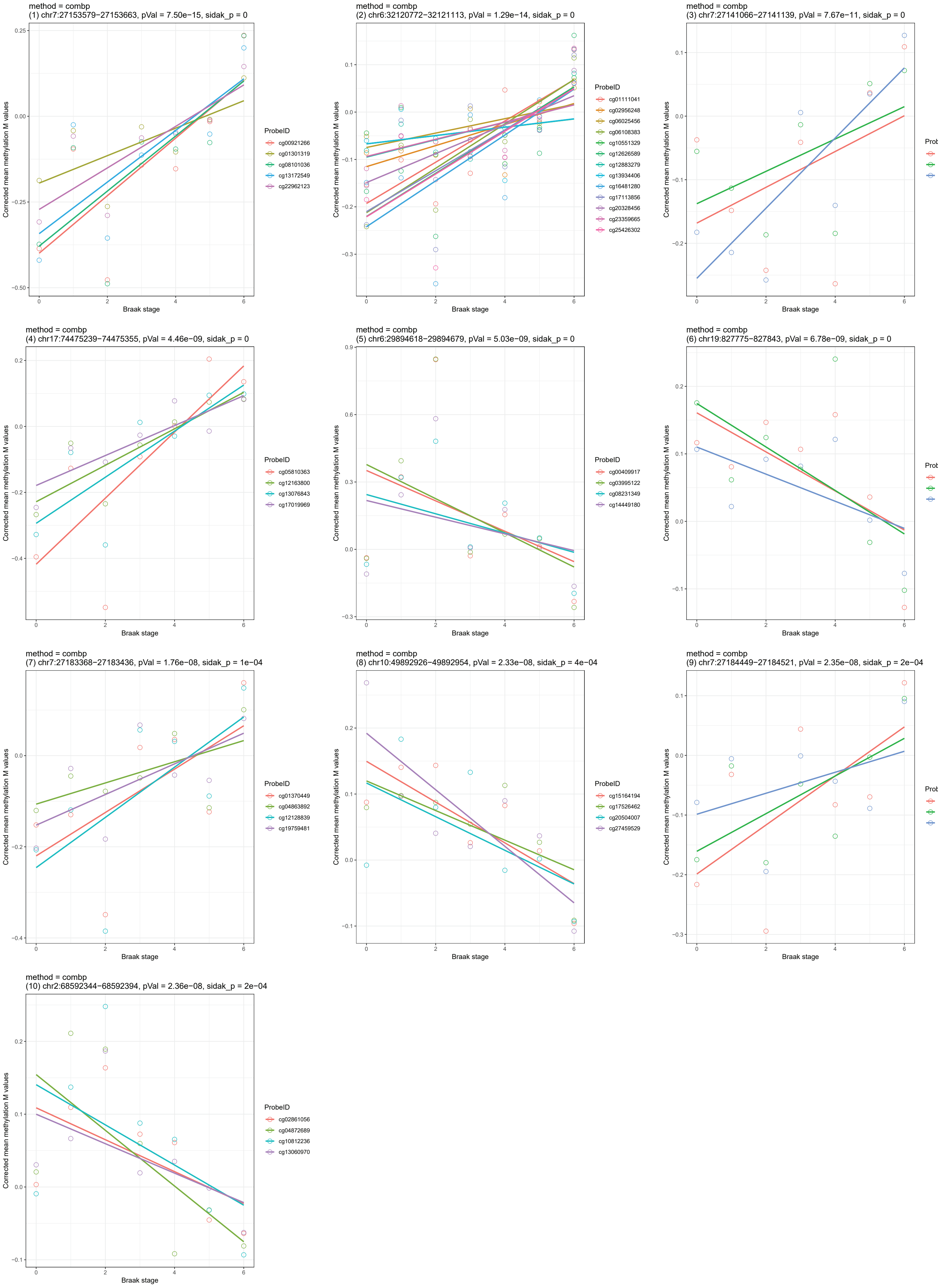
method = combp
(1) chr7:27153579−27153663, pVal = 7.50e−15, sidak_p = 0
ProbeID
cg00921266
cg01301319
cg08101036
cg13172549
cg22962123
method = combp
(2) chr6:32120772−32121113, pVal = 1.29e−14, sidak_p = 0
ProbeID
cg01111041
cg02956248
cg06025456
cg06108383
cg10551329
cg12626589
cg12883279
cg13934406
cg16481280
cg17113856
cg20328456
cg23359665
cg25426302
method = combp
(3) chr7:27141066−27141139, pVal = 7.67e−11, sidak_p = 0
ProbeID
cg02803819
cg13661519
cg21464565
method = combp
(4) chr17:74475239−74475355, pVal = 4.46e−09, sidak_p = 0
ProbeID
cg05810363
cg12163800
cg13076843
cg17019969
method = combp
(5) chr6:29894618−29894679, pVal = 5.03e−09, sidak_p = 0
ProbeID
cg00409917
cg03995122
cg08231349
cg14449180
method = combp
(6) chr19:827775−827843, pVal = 6.78e−09, sidak_p = 0
ProbeID
cg10752406
cg15610437
cg16643542
method = combp
(7) chr7:27183368−27183436, pVal = 1.76e−08, sidak_p = 1e−04
ProbeID
cg01370449
cg04863892
cg12128839
cg19759481
method = combp
(8) chr10:49892926−49892954, pVal = 2.33e−08, sidak_p = 4e−04
ProbeID
cg15164194
cg17526462
cg20504007
cg27459529
method = combp
(9) chr7:27184449−27184521, pVal = 2.35e−08, sidak_p = 2e−04
ProbeID
cg02106682
cg03368099
cg07049592
method = combp
(10) chr2:68592344−68592394, pVal = 2.36e−08, sidak_p = 2e−04
ProbeID
cg02861056
cg04872689
cg10812236
cg13060970
Corrected mean methylation M values
Braak stage
